# Gut-Brain Hydraulics: Brain motion and CSF circulation is driven by mechanical coupling with the abdomen

**DOI:** 10.1101/2025.01.30.635779

**Authors:** C. Spencer Garborg, Beatrice Ghitti, Qingguang Zhang, Joseph M. Ricotta, Noah Frank, Sara J. Mueller, Denver I. Greenawalt, Kevin L. Turner, Ravi T. Kedarasetti, Marceline Mostafa, Hyunseok Lee, Francesco Costanzo, Patrick J. Drew

**Affiliations:** Penn State Neuroscience Institute – University Park, The Pennsylvania State University, University Park, PA 16802; Center for Neural Engineering, The Pennsylvania State University, University Park, PA 16802; Department of Biomedical Engineering, The Pennsylvania State University, University Park, PA 16802; Department of Engineering Science and Mechanics, The Pennsylvania State University, University Park, PA 16802; Auckland Bioengineering Institute, The University of Auckland, Auckland, New Zealand; Department of Physiology, Michigan State University, East Lansing MI; Department of Mechanical Engineering, The Pennsylvania State University, University Park, PA 16802; Center for Quantitative Imaging, The Pennsylvania State University, University Park, PA 16802; Department of Biology, The Pennsylvania State University, University Park, PA 16802; Department of Mathematics, The Pennsylvania State University, University Park, PA 16802; Department of Neurosurgery, The Pennsylvania State University, University Park, PA 16802

**Author notes:** **Corresponding author:** Patrick J. Drew.

## Abstract

The brain moves within the skull, but the drivers and function of this motion are not understood. We visualized brain motion relative to the skull in awake head-fixed mice using high-speed, multi-plane two-photon microscopy. Brain motion was primarily rostrally and laterally directed, and was tightly correlated with locomotion, but not with respiration or the cardiac cycle. Electromyography recordings in abdominal muscles and microCT reconstructions of the trunk and spinal vasculature showed that brain motion was driven by abdominal muscle contractions that activate a hydraulic-like vascular connection between the nervous system and the abdominal cavity. Externally-applied abdominal pressure generated brain motion similar to those seen during abdominal muscle contractions. Simulations showed that brain motion drives substantial volumes of interstitial fluid through and out of the brain (at volumetric rates several times higher than production) into the subarachnoid space, in the opposite direction of fluid flow seen during sleep. The brain is hydraulically linked to the abdominal compartment, and fluid flow in the brain is coupled to body movements, providing a mechanism by which the mechanics of the viscera directly impact brain health.

## Introduction

Brain motion is a ubiquitous, but poorly investigated phenomenon ^1,2^. In anesthetized animals, brain motion is closely tied to cardiac pulsations and respiration ^3^, but in unanesthetized animals, brain motion is usually typically associated with locomotion and other body movements ^2,4^. In mice, brain motion observed with two-photon microscopy is on the order of a few microns ^2,5,6^ and is primarily within the imaging plane (medial-lateral/rostral-caudal).

Despite the ubiquity of brain motion in the awake animal, its origins are not well understood. A force must be exerted on the brain for it to move, but the central nervous system has been considered to be largely mechanically insulated from the rest of the body by the skull and vertebrae. Despite this partitioning, during locomotion the intracranial pressure (ICP) of mice rises from a baseline of approximately 5 mmHg to more than 20 mmHg ^7,8^, indicating that substantial mechanical forces are rapidly applied to the brain during body movements. The increase in ICP during locomotion is not due to dilation of blood vessels within the brain, as the hemodynamic response lags both the pressure increase and the onset of locomotion by approximately one second^9,10^. Furthermore, maximally dilating the vessels of the brain does not increase ICP nearly as much as locomotion ^7^. These pressure changes are unlikely to be simply an epiphenomenon because brain motion during locomotion excites sensory neurons in the dura ^11^, indicating that the motion of the brain is actively monitored and may serve a physiological role.

One potential physiological purpose for brain motion is to circulate interstitial fluid (ISF) and cerebrospinal fluid (CSF) in the brain. As the brain lacks a lymphatic system to remove waste, it depends on mechanical forces exerted on it by pulsation ^12^ and dilation and constrictions ^13-15^ of arteries to help circulate fluid though the glymphatic system. During sleep, CSF is driven into the brain along the periarterial spaces of penetrating arteries by slow, alternating dilation and constriction of the vessel ^16-19^. The patterns of fluid flow in the brain are markedly different in the awake animal, where tracers do not enter the cortex ^20^, though the reasons for this difference between sleep and wake CSF flow is not completely understood. The large forces that drive brain motion are also likely to drive movement of CSF, potentially in very different patterns than those that are seen during sleep. However, understanding these fluid flows requires a detailed characterization of the mechanical dynamics of the brain.

We used high-speed, multiplane two-photon microscopy to measure motion of the dorsal cortex in awake head-fixed mice. Brain motion relative to the skull was highly correlated with locomotion, and primarily in the rostral and lateral directions. Using microcomputed tomography (microCT), we visualized the vertebral venous plexus (VVP), a network of valveless veins that connect the abdominal cavity to the spinal cavity. This vascular network is a hydraulic system that transmits pressure from the abdomen to the spinal cavity, where it impacts the central nervous system ^21^. We found that brain movements closely followed the contraction of abdominal muscles, and that passive pressure to the abdomen in anesthetized mice could recapitulate the rostral brain shift within the skull that was seen in the awake mouse. To reveal motion-induced fluid flow inside and around the brain, we performed poroelastic brain tissue simulations constrained by our measurements of brain motion and known intracranial pressure changes. In these simulations, brain motion drove movement of substantial amounts of fluid (several times the amount of CSF is produced in a comparable time) within the brain out into the subarachnoid space, the opposite direction of fluid flow seen during sleep. Our work demonstrates that the brain is mechanically linked to the abdomen and that this connection is a novel and important driver of fluid flow in the awake brain.

## Results

We used two-photon microscopy to quantify brain motion relative to the skull in 24 Swiss Webster mice (12 male) that were head-fixed upon a spherical treadmill. We simultaneously imaged brain cells expressing green fluorescent protein ^22^ and fluorescent microspheres attached to a polished and reinforced thinned-skull (PoRTS) window ^23^. This was accomplished by integrating an electrically tunable lens behind the microscope objective to rapidly (39.55 frames/sec, 19.78 frames/sec per plane) alternate between two focal planes on the skull surface and in the brain (Fig 1, SFig 1,2), separated by ~90µm. Tracking of microspheres showed that skull movement was usually less than 1µm (SFig 3), demonstrating the stability of the head fixation apparatus and that the displacement perpendicular to the imaging plane was minimal relative to the size of point spread function in z (SFig 2c). The motion of the brain relative to the skull was primarily in the rostral and lateral directions (Fig 1e) and was strongly correlated with locomotion (Fig 2d, SFig 4a,b). We found uniform displacements across the field of view (SFig 5, Mov 1), indicating that there is minimal strain over the imaged area and that displacement can be captured with rigid translation.

**Figure 1.**
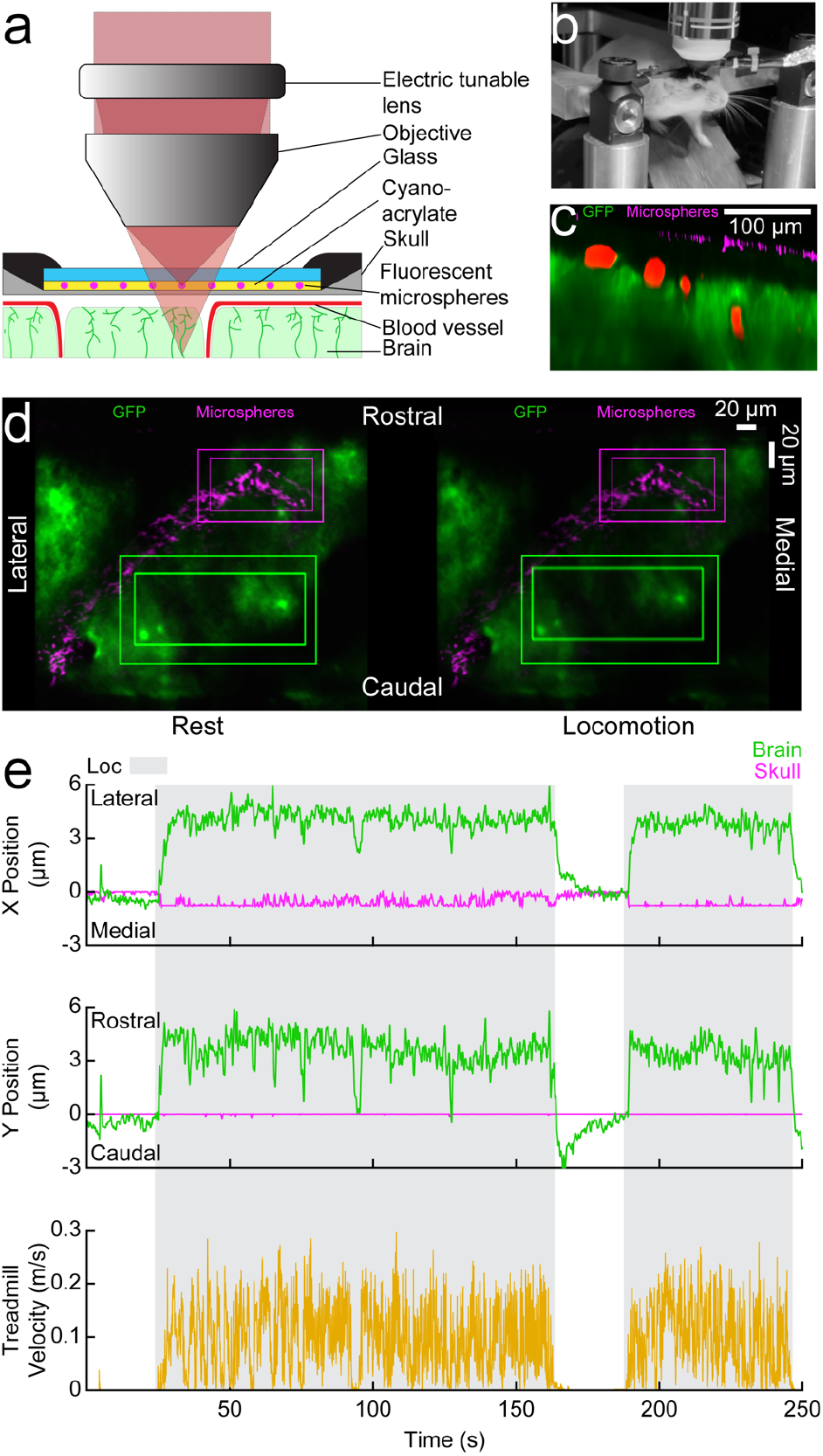
Two-photon imaging of brain motion relative to the skull. **a**. Rapid changes in the curvature of the fluid-filled lens move the focal point between the brain and the fluorescent microspheres adhered to the surface of the thinned skull. **b**. Head-fixed mouse on a treadmill. **c**. A representative X-Z image through a typical thinned-skull window. The GFP-expressing brain (green) and fluorescent microspheres (magenta) on the thinned skull are separated by the subarachnoid space. **d**. Images of the brain (green) and microspheres (magenta) during a stationary period (left) and locomotion (right). The outer bounding boxes enclose the search area for the template-matching algorithm, while the inner bounding boxes represent the target used to track movement. There is a rostro-lateral shift of the brain during locomotion when compared to the rest image (visible in the displacement of the inner box) while the skull remains in its resting position. **e**. An example of measured brain motion. Locomotion events, shown in gray, drive rostro-lateral motion of the brain (green) while the skull (magenta) remains stationary.

### The brain motion is primarily in the rostral direction and is linked to locomotion

To quantify patterns in the direction of motion, we imaged brain motion during locomotion from 134 sites in frontal, somatosensory and visual cortex and performed principal component analysis on the brain displacement (Fig 2). The magnitude of each displacement vector was determined by averaging the largest 20% of the displacements from the baseline origin (Fig 2a). We observed that the motion of the brain during locomotion was primarily in the rostral and lateral directions relative to the resting baseline position (Fig 2b, Mov 2,3,4). Brain motion amplitude was larger in males than in females (SFig 6). When we looked at the power spectrum of the motion, we observed the motion was primarily at low (<0.1 Hz) frequencies (Fig 2c), and it was strongly correlated to locomotion in both directions (Fig 2d, SFig 4a,b). We did not observe any appreciable brain movement at respiration or heart rate frequencies (Fig 2c) in awake mice. However, respiration-induced movement was detected under deep isoflurane anesthesia (SFig 7, Mov 5).

**Figure 2.**
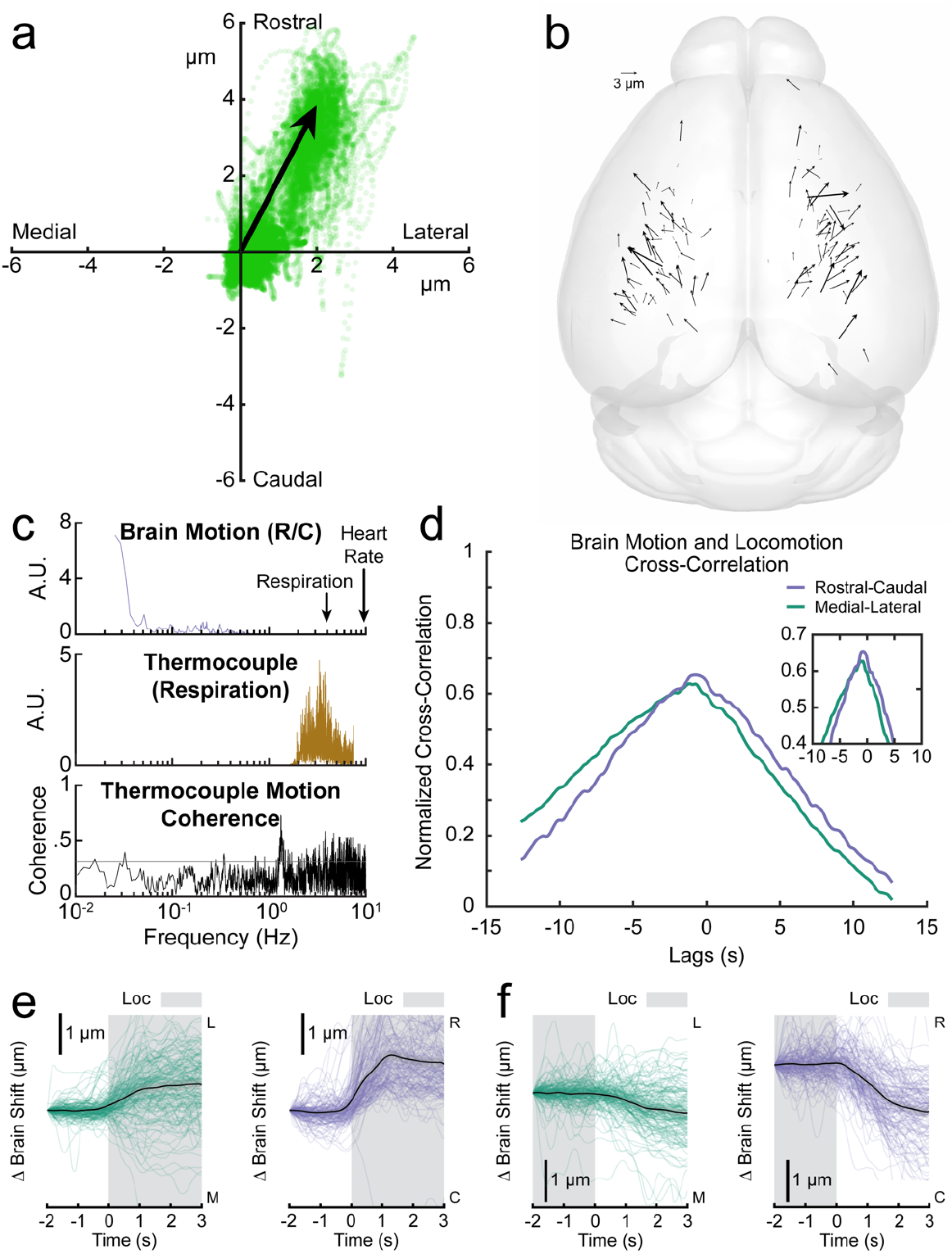
The brain moves rostrally and laterally within the skull in locomoting mice. **a**. The net displacement of the brain in each frame (from data in Fig 3d) plotted as a x-y scatterplot. The displacement vector is taken to be the first principal component of the data, and the magnitude is calculated as the mean of the 80^th^ to 100^th^ percentile of the displacement magnitudes. **b**. A plot of displacement vectors for different imaging locations on the brain (N=134 sites in 24 mice). There is a noticeable rostro-lateral brain movement trend in both hemispheres. **c**. Power spectrums of rostral-caudal brain motion (top) and respiration (middle), showing there is no appreciable brain motion at the respiration frequency. Plotted at the bottom is the coherence between rostral-caudal brain motion and respiration. A lack of overlap in the frequency components of the signals and a low coherence between them (confidence = 0.319) suggest that the observed motion is not driven by respiration or heartbeat. **d**. Cross-correlations between the brain motion and locomotion signals from (Fig 3d). **e**. Locomotion-triggered rostral-caudal and medial-lateral brain motion. Each colored line represents the locomotion-triggered average for a single trial and the black line is the mean with the shading showing the 90 percent confidence interval. The brain begins to move rostrally and laterally slightly prior to locomotion. **f**. Triggered averages of the cessation of locomotion. The brain moves caudally and medially to return to baseline following the transition from locomotion to rest.

The skull and brain are separated by the dura, a vascularized membrane surrounding the subarachnoid space ^24,25^. In one instance, we were able to simultaneously record movement of dural vessels labeled with green fluorescent proteins, microspheres on the skull, and the brain. This allowed us to determine if the dura motion more closely resembled brain or skull movement during locomotion. We performed tracking on the three focal planes separately (Mov 4) and observed that the dura had similar dynamics to the skull. We generated locomotion triggered averages of brain motion and found a close relationship between locomotion and movement of the brain (Fig 2e), though the motion of the brain in many cases started prior to locomotion onset.

These results demonstrate that in awake mice, locomotion is linked to brain motion while respiration and heart rate are not substantial contributors to brain motion. However, brain motion frequently preceded the onset of locomotion, suggesting that locomotion in and of itself does not cause brain motion within the skull.

### Brain motion follows abdominal muscle contractions

The brain motion we observed often slightly preceded locomotion, which indicated that a force was being applied to the brain prior to locomotion onset. Intracranial pressure (ICP) in mice increases sharply during locomotion (from 5-10mmHg to >20 mmHg) ^7^, indicating that there are large forces at work on the brain. The increase in ICP also precedes the onset of locomotion, and this cannot be attributed to vasodilation as it lags locomotion ^26^. Furthermore, brain motion is unlikely to be due to postural changes as these also lag locomotion onset. ^27^We hypothesized that abdominal muscle contractions might contribute to brain motion because movements are preceded by abdominal muscle activation to stiffen the core in anticipation of body motion. We implanted electromyography (EMG) electrodes in the abdominal muscles of 24 mice while simultaneously monitoring brain movement (Fig 3a). EMG power increased prior to the onset of locomotion (Fig 3b,c), and there was a strong correlation between EMG power, which tracks muscle tension, and the motion of the brain (Fig 3f, SFig 6c,d). When we aligned brain motion to the onset of locomotion and to the onset of EMG activity, we observed that the motion invariably lagged EMG activity (Fig 3g,h, SFig 8), but often preceded locomotion, which suggested that abdominal muscle contraction prior to locomotion drove the displacement of the brain.

**Figure 3.**
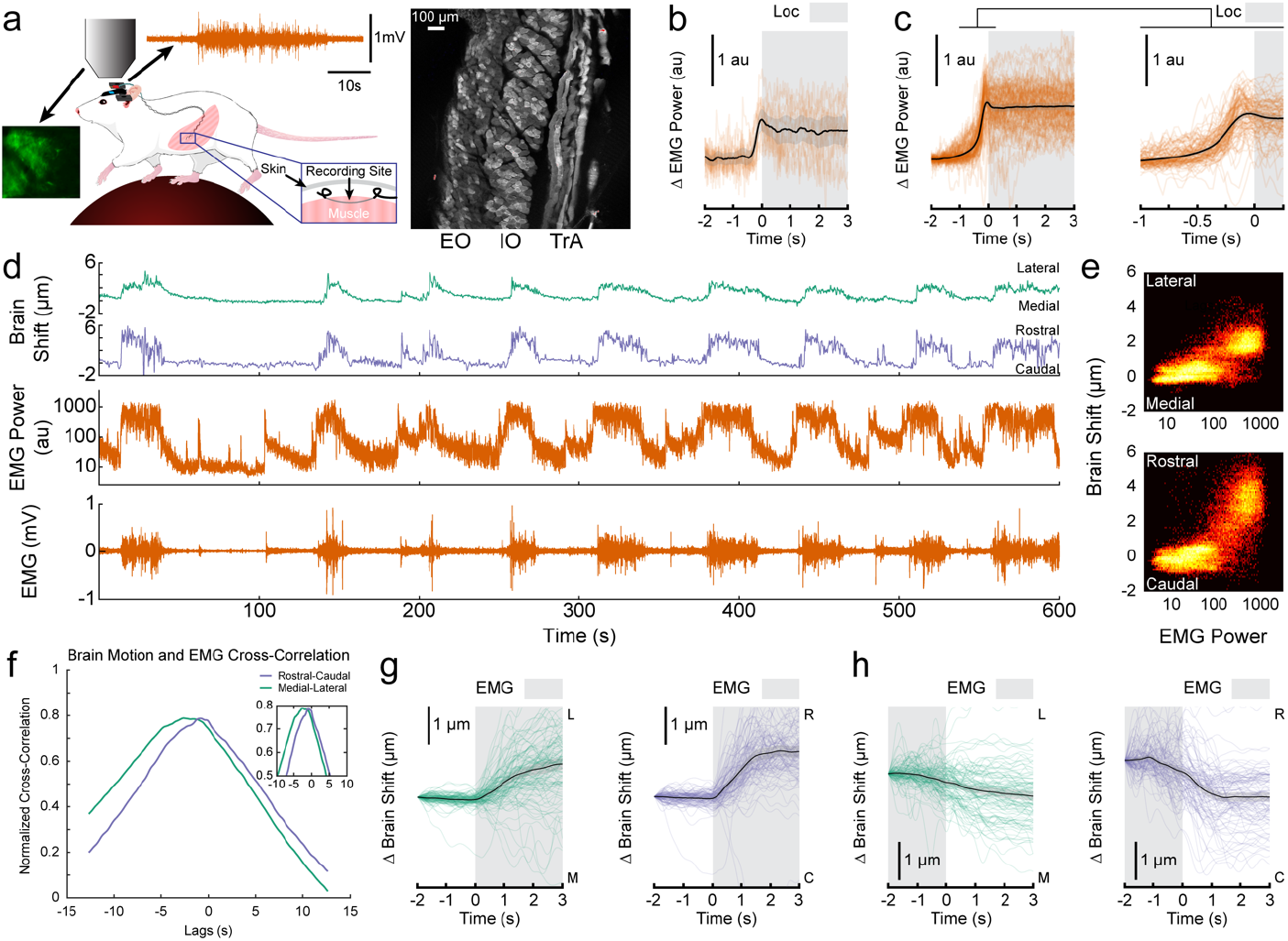
Abdominal muscle activation predicts brain motion. **a**. EMG electrodes were implanted in the abdominal muscles (left), which consist of three layers (right). **b**. The locomotion-triggered abdominal EMG power (orange) from a single trial representative trial (data in **d**). Black line denotes mean, shading the 90 percent confidence interval. **c**. The locomotion-triggered abdominal EMG averages for all trials (orange). The expanded view around the trigger (right) shows that the abdominal EMG increases prior to the onset of locomotion. **d**. Representative brain displacement and abdominal EMG. Note the degree of correlation between abdominal muscle contraction and motion of the brain within the skull. **e**. Two-dimensional histograms of abdominal EMG power and brain displacement in a single trial (data in **d**). **f**. Cross-correlation between abdominal muscle EMG power and brain position for data in **d. g**. EMG-triggered averages for rostral-caudal and medial-lateral brain motion. Each colored line represents the EMG-triggered average for a single trial and the black line represents the mean with a 90 percent confidence interval. The brain begins to move rostrally and laterally simultaneously with the onset of abdominal muscle activation. **H**. Triggered averages of the cessation of abdominal muscle activity. The brain moves caudally and medially to return to baseline around the time that the abdominal muscles relax.

We then tested the relationship between brain motion and recruitment of abdominal musculature in non-locomotor regimes. Respiration conditionally recruits abdominal musculature: While exhalation does not recruit abdominal musculature at rest, respiratory distress conditionally elicits active expiration through abdominal muscle contraction^28^. Under deep anesthesia, we observed active expiration as revealed by the onset of abdominal EMG power bursts locked to respiratory rhythm. These EMG bursts were also locked to brain motion (SFig 7b,d, Mov 5). During periods of shallow, rapid breathing, both EMG power and brain motion were reduced (SFig 7d). Finally, we observed instances of abdominal muscle activation and brain motion in the absence of locomotion (SFig 7e, Mov 3). These results show that across a wide variety of physiological regimes, abdominal muscle activation is responsible for driving brain motion.

### Vertebral venous plexus provides a hydraulic link between abdomen and CNS

How could forces generated by abdominal muscle contraction reach the brain? In humans, abdominal muscle activation drives an increase in intra-abdominal pressure (IAP) ^29^. These increases in IAP are communicated to the brain and spine ^30^ via the vertebral venous plexus (VVP)^31^, a network of valveless veins that connect the abdomen and spinal canal ^32^. The VVP is thought to function like a hydraulic system that provides circulatory regulation during postural changes, in which pressure in one compartment (the abdomen) exerts pressure on another (the spinal column) via the movement of fluid (blood) from higher-pressure regions to lower-pressure regions. However, whether mice possess a functional VVP was unknown. We filled the vascular system of a mouse with a radiopaque tracer, imaged it using microCT, and reconstructed the vasculature around the vertebral column (Fig 4, SFig 9, Mov 6).

**Figure 4.**
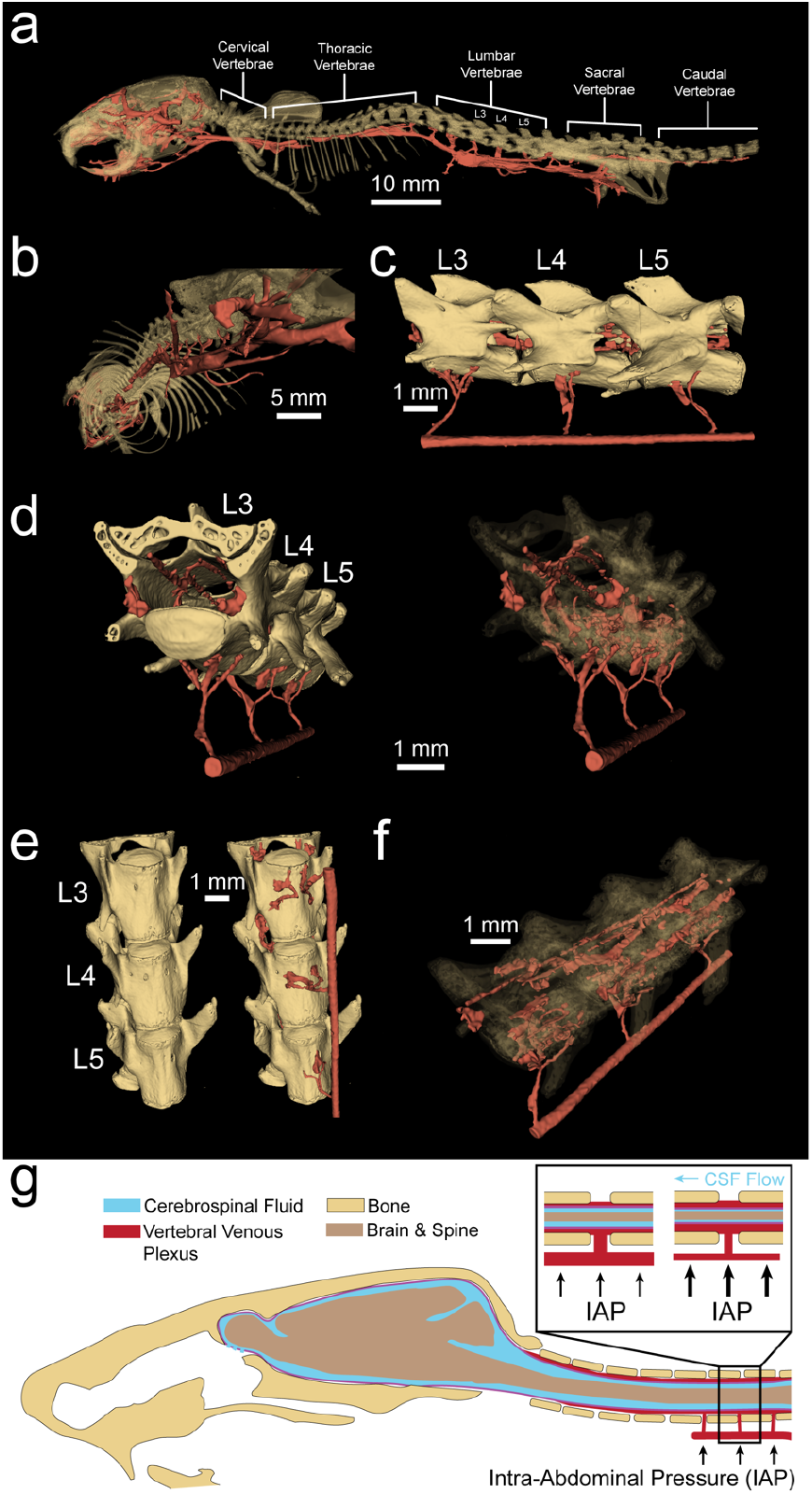
The vertebral venous plexus (VVP) provides a mechanism for abdominal pressure changes to influence brain motion. **a**. Segmented microCT scan of a mouse skeleton (gold) and vasculature (red). **b**. Venous connections from the caudal vena cava are shown to bifurcate prior to entering the lumbar vertebrae. **c**. Connections from the caudal vena cava inferior to the L3, L4, and L5 vertebrae penetrate the vertebrae and connect to vasculature surrounding the spinal cord. **d**. Veins run longitudinally along the interior of the vertebrae (left). The venous bifurcations connect the caudal vena cava and vasculature within the spine. **e**. Small holes in the ventral surfaces of the lumbar vertebrae provide an entrance for the venous projections to connect to vasculature surrounding the dural sac within the column. **f**. A semi-transparent view of the vertebrae provides a complete look at the caudal vena cava, the vessels that run the length of the vertebral interior, and the connections between them. **g**. Increased intrabdominal pressure forces blood from the caudal vena cava to the VVP within the vertebral column. The increased blood volume in an enclosed space applies pressure to the dural sac, forcing the cranial CSF flow that generates brain motion.

We found the lumbar and sacral vertebra, but not the thoracic vertebrae, had small ventral foramina that communicate with the spinal canal (Fig 4e). These foramina were typically in pairs and located on both sides of the vertebral body, though some vertebrae possess only one. Blood vessels were observed to clearly communicate through these holes into a vascular network that lined the walls of the spinal cavity, providing a physical link between the abdominal compartment and the CNS. The diaphragm partitions the thoracic and abdominal cavities while also separating the VVP-connected lumbar and sacral vertebrae from the thoracic vertebrae that lack VVP communication pathways. This separation allows the VVP to transmit abdominal (but not thoracic) pressure changes to the CNS. In humans, intrabdominal pressures rise drastically (~90mmHg) when the abdominal muscles are contracted ^27^. A pressure increase of this magnitude will drive some of the blood in the abdomen into the spinal canal, narrowing the dural sac. This results in cranial CSF flow that raises ICP and drives brain motion (Fig 4f, Ani 1).

### Brain motion induced by externally-applied abdominal pressure

If the mechanical coupling between the abdomen and central nervous system via the VVP drives brain motion, then we reasoned that passively applied pressures to the abdomen should drive similar brain movements. To test this idea, we constructed a pneumatic pressure cuff (SFig 10) to apply controlled pressure to the abdomen of lightly anesthetized (~1% isoflurane in oxygen) mice (Fig 5). We observed that the brain began moving rostrally and sometimes laterally within the skull shortly following the onset of the abdominal compression (Fig 5e, Mov 7). Furthermore, the brain began moving back to its baseline position immediately upon relief of the abdominal pressure. This suggests that abdominal pressure can rapidly and significantly alter the position of the brain within the skull.

**Figure 5.**
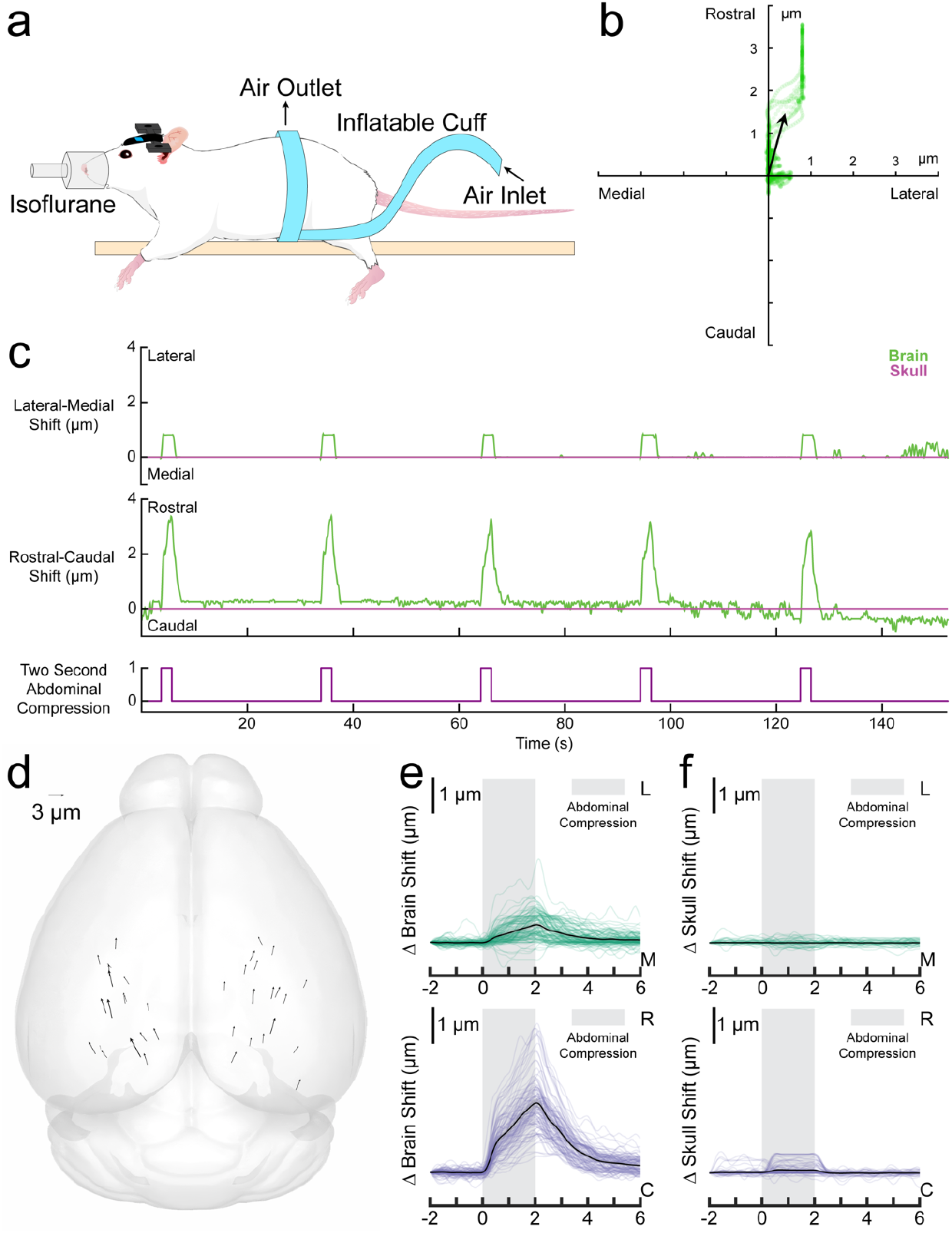
Pressure applied to the abdomen of anesthetized mice resulted in rostro-lateral brain motion. **a**. The mouse was lightly anesthetized with isoflurane and wrapped with an inflatable belt. **b**. Displacement of the brain relative to the skull (green) for a single abdominal compression trial (data in **c**). The brain was displaced rostrally and slightly laterally. **c**. Displacements of the brain (green) and skull (magenta) during abdominal compressions delivered to the anesthetized mouse (blue). **d**. Brain displacement during abdominal compression trials across the brain (36 locations in 6 mice). The motion trend is in the rostro-lateral direction, as seen with brain motion during locomotion. Generated using brainrender^58^. **e**. Abdominal compression-triggered average of brain motion for each trial in the medial-lateral (green) and rostral-caudal (blue) direction. The black line shows the mean, shading the 90 percent confidence interval. The brain begins moving immediately upon abdominal pressure application and continues to displace as the compression continues. Upon pressure release, the brain quickly returns to baseline. **f**. Abdominal compression-triggered skull motion averages for each trial in the medial-lateral (green) and rostral-caudal (blue) direction.

### Simulations show motion generates fluid flow out of the brain

The movement of CSF/ISF into, through, and out of the brain through the glymphatic system is important for the clearance of waste ^12^, and recent work has pointed to the mechanical forces generated by the dilation or constrictions of blood vessels in generating this fluid motion ^13-15,33^. We hypothesized that the large movements that we see of the entire brain could drive fluid motion of a different sort. However, while fluid flow in the subarachnoid space and ventricles can be visualized in certain instances ^17,34^, the rapid dynamics of any motion-driven fluid flow through the parenchyma and around the brain in the awake animal is not accessible to current imaging techniques in behaving mice. Therefore, we simulated the fluid flow produced by a squeezing action of the spinal cord using a poroelastic model of the brain and spinal cord (Fig 6). Our axisymmetric model of a brain with simplified geometry incorporated a rostral outflow point corresponding to the cribriform plate, and a compliant vascular portion in the brain corresponding to the bridging veins ^35^ to buffer pressure changes (Fig 6a). We simulated pressure application to the distal spinal cord to mimic abdominal muscle contraction such that the model gave ICP changes and brain motion consistent with our experimental observations (Fig 6b,c). We then used the model to see what the corresponding fluid flows (Fig 6d,e) were in and around the brain. Surprisingly, there was a net flow of fluid *out* of the brain (Fig 6e), into the subarachnoid space. The direction of the fluid flow relative to the solid motion can be deduced from the streamlines of the filtration velocity (Fig 6d). This brain motion induced flux was large, corresponding to approximately five times the normal CSF production rate ^36^ (Fig 6d), meaning that brain-motion-induced fluxes should be the dominant driver of fluid flow in the awake brain. Intriguingly, these flows are in the opposite direction of the glymphatic flow seen during sleep ^19^ and consistent with experimental observations that tracers infused into the cisterna magna in awake mice do not enter in to the cortex ^20^. Our simulations showed that flows across the cranial and spinal SAS are orders of magnitude larger than those across the ventricle and central canal surfaces (Fig 6e). Additionally, quantitative details about fluid flows within the brain and SAS domains can be found in SFig 11a-c.

**Figure 6.**
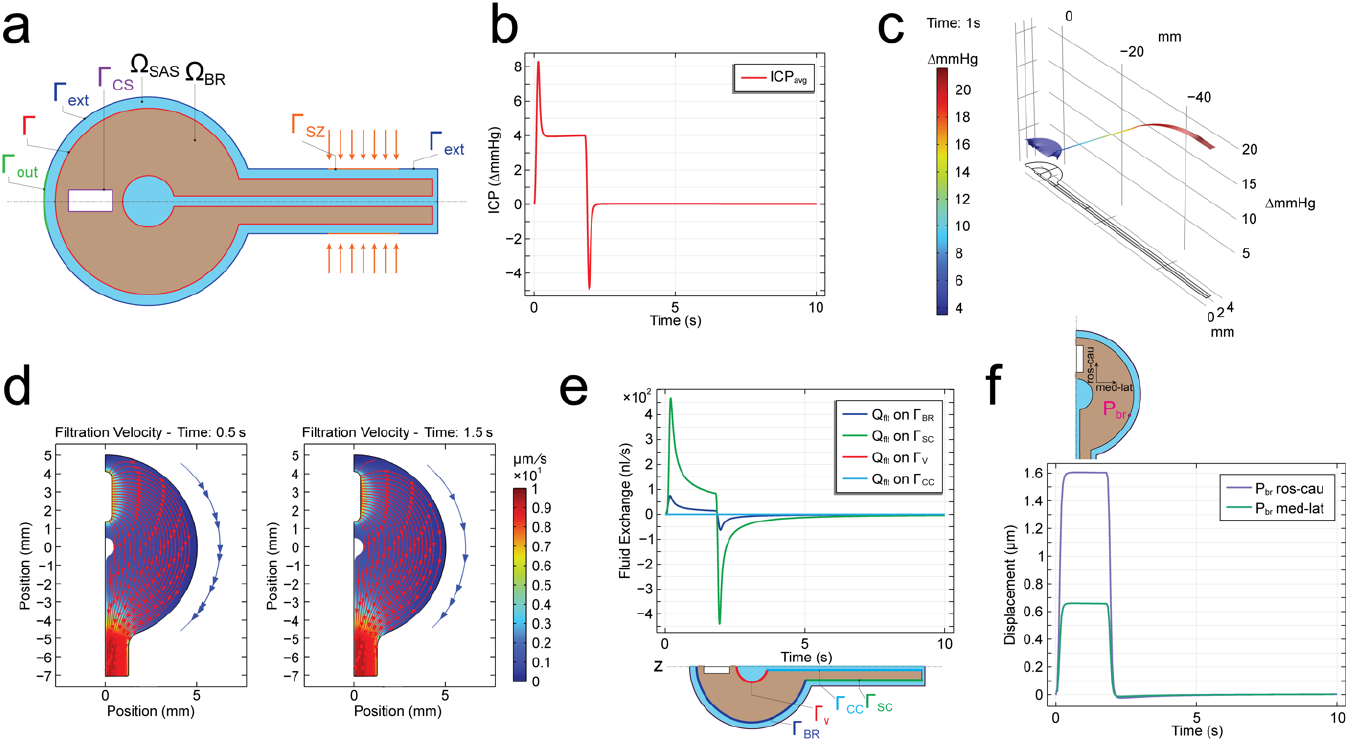
Computational results from a finite element simulation of the flow induced by a squeeze of intensity *p*_0_ = 20mmHg applied over the SZ. The duration of the squeeze pulse is 2s. The duration of the simulation is 10s. The simulation is based on Equations (1)—(9). The boundary conditions are described in the Supplementary Material. The parameters used in the simulation are found in Supplementary Table 1. **Note: the resistance scaling factors adopted here are** α_cs_ = 10^6^ **and** α_out_ = 6 × 10^8^. **a**. Initial geometry (not to scale) detailing model domains and boundaries. Ω_BR_: brain and spinal cord domain (pale pink); Ω_SAS_: CSF-filled domain (cyan); Γ: Ω_BR_ − Ω_SAS_ interface (red); Γ_ext_: external boundary of meningeal layer (blue); Γ_sZ_: squeeze zone (orange); Γ_out_: outlet boundary representing the cribriform plate CSF outflow pathway (green); Γ_cs_: central sinus boundary (purple). **b**. Average of pore pressure (in mmHg) over Ω_BR_ excluding the spinal cord over time. **c**. Spatial distribution of pore pressure (in mmHg) over Ω_BR_ ∪ Ω_SAS_ at *t* = 1 s during the squeeze pulse. **d**. Streamlines of filtration velocity ***ν***_flt_ (i.e., curves tangent to filtration velocity field; red arrows) within Ω_BR_ excluding the spinal cord, at *t* = 0.5 s (left) and *t* = 1.5 s (right) during the squeeze pulse, overlaying the color plot of the filtration velocity magnitude (in μm/s), computed as 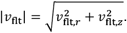. Because the SAS is extremely thin, it is not meaningful to show a full plot of the streamlines in the SAS. This said, the blue line with arrows placed on the right side of each streamline plot is meant to indicate the direction of flow in the SAS at the corresponding time. **e**. Volumetric fluid exchange rate Q_flt_ (in nL/s) over time across: the brain shell surface Γ_bζ_ (blue), spinal cord surface Γ_sc_ (green), ventricle surface Γ_***x***_ (red), and central canal surface Γ_cc_ (light blue). Q_flt_ > 0: fluid flow from Ω_BR_ into Ω_SAS_. Q_flt_ is computed as the integral of the normal component of filtration velocity over the surfaces indicated. The plot displays 4 lines, two that are easily seen (blue and green lines), and two that overlap and appear as horizontal lines near zero (red and light blue lines). This is due to the different orders of magnitude of Q_flt_ across the different portions of Γ. **f**. Rostro-caudal (blue) and medio-lateral (green) motion of point *P*_br_ on the brain surface (shown in the inset) over time caused by the squeeze pulse.

We saw similar patterns of fluid flow out of the brain when we varied the outflow resistance/bridging vein compliance within ranges that produced physiologically realistic ICP changes and brain motions, suggesting that these results hold generally (SFig 12,13). Finally, the simulations predicted rostral/medial motion at the rostral tip of the brain (Fig 6f, SFig 11d). We performed imaging of brain motion dynamics in the corresponding position in the brain, the olfactory bulb, and also saw rostral/medial motion (SFig 14, Mov 8), indicating that our simple model geometry is capturing the fundamental aspects of brain motion. In toto, these simulations show that brain motion causes large fluid flows out of the brain, in the opposite direction of glymphatic flow during sleep, potentially explaining why the quiescence during sleep is required to drive fluid flow through the glymphatic system.

The parameter values used in the simulation discussed herein were the ones that allowed us to obtain some agreement with two essential empirical measurements carried out in the study, namely brain surface displacement and intracranial pressure. In the Supplementary Material, we performed simulations adopting different values of the resistance at the outlet and offered by the central sinus (SFig 12,13). In both cases, these resistances play an important role in achieving the observed values of intracranial pressure.

## Discussion

Our work shows that the brain is not mechanically isolated from the body, but rather is very closely coupled to the abdominal cavity via the VVP. The effect on fluid flow by motion of the brain could help explain why injected tracers do not enter into the cortex in awake animals but do so readily during sleep ^20^. In humans, the VVP is thought to help buffer ICP ^31^, but its role in rodents is puzzling since the hydrostatic pressure gradients in a mouse will be much smaller than those in a human, both overall and relative to their respective arterial pressures. This hydraulic system can generate brain motion within the skull and drive CSF flow out of the brain into the subarachnoid space. Tension by spinal nerves^37^ during the motor act of locomoting are unlikely to have generated brain motion in this experiment because we observed brain motion in the absence of changes in body configuration (SFig 7e, Mov 3). In fact, our simulations predicted that brain motion is induced by the force exerted by the VVP on the spinal cord (SFig 11f).

One caveat is that the mice were head fixed, preventing the normal forces generated by head motion from acting on the brain. However, the forces created by head movement in mice are much smaller than those generated by IAP and ICP changes. Measurements in freely behaving mice show self-generated accelerations of order 1g ^38^, resulting in a force of ~4 millinewtons (9.8m/s^2^*0.4g brain mass). The forces generated by a 10 mmHg anterior-posterior pressure change^7^ on the ~30 mm^2^ coronal cross-sectional area of the mouse brain will be substantially larger than those generated by head motion, on the order of ~40 millinewtons (1333N/m^2^ * 30×10^-6^m^2^). In contrast, head motion-generated forces will be greater in humans where the brain mass is several orders of magnitude larger, though ICP changes are also greater in humans than in mice ^39^.

Our results also demonstrate a novel and immediate link between the brain and viscera state, mediated by abdominal pressure. Obesity ^40^ elevates IAP, which could disrupt the normal flow of blood between the abdominal cavity and spinal canal and/or lead to remodeling of the VVP. Alteration of blood flow and pressure gradients between the abdomen and spinal canal could reduce the movement of the brain and CSF circulation, contributing to the adverse effects of obesity on cognitive function ^41^. Reduction of abdominal pressure though voiding or defecation ^42^ may partly contribute to their impacts on cognition ^43^. Mechanical coupling between the abdomen and the brain is especially interesting considering the functional mechanosensitive channels in CNS neurons ^44^ and glia ^45^, as the forces that cause brain motion could also activate mechanosensitive channels in the brain. In addition to interoceptive pathways in the viscera, the direct signaling through mechanical forces to the brain may play a role in communicating internal states to the brain.

The simulations also indicate the importance of accounting for the deformation of vascular compartments, such as the central sinus. This observation adds to considerations coming from existing literature on the glymphatic system, emphasizing the importance of capturing the interaction between vascular dynamics and brain motion in the understanding of brain waste clearance.

## Methods

All experiments were done with the approval of the PSU Institutional Animal Care and Use Committee. We imaged 30 (15 male) Swiss Webster (Charles River, #024CFW) mice. We chose Swiss Webster mice as the dorsal skull is substantially flatter than other mouse strains, their skull bones are fused, and their larger size made it easier to implant abdominal muscle EMG electrodes.

One month prior to window implantation, expression of GFP across brain cells^22^ was induced using retroorbital injection of 10 μL AAV (Addgene #37825-PHPeB, 1×10^13^ vg/mL) in 90 μL H_2_O (SFig 1b). We implanted a PoRTS window, with the additional step that fluorescent microspheres were applied to the surface of the skull (Fig 1c, SFig 1a). In all mice, EMG electrodes were implanted in the abdominal muscles. Mice were then habituated to head fixation over several days before imaging.

### Window and abdominal EMG surgery

Mice were anesthetized with isoflurane (5% induction, 2% maintenance) in oxygen throughout the surgical procedure. The scalp was shaved, and an incision was made from just rostral of the olfactory bulbs to the neck muscles, which was opened to expose the skull. A custom 1.65mm thick titanium head bar was adhered to the skull using cyanoacrylate glue (Vibra-Tite, 32402) and dental cement. To assist with head bar stabilization, two small self-tapping screws (J.I. Morris, F000CE094) were inserted in the frontal bone without penetrating the subarachnoid space and were connected to the head bar with dental cement. A PoRTS window was then created over both hemispheres ^23^. Windows typically spanned an area from lambda to rostral of bregma and were up to 0.5 cm wide, spanning across somatosensory and visual cortex. This allowed for maximum viewable brain surface. The skull was thinned and polished, and 1-μm diameter fluorescent microspheres (Invitrogen, T7282) were spread across the surface of the thinned-skull areas and allowed to dry. They were then covered with cyanoacrylate glue and a 0.1-mm thick borosilicate glass piece (Electrode Microscopy Sciences, 72198) cut to the size of the window. The position of bregma was marked with a fluorescent marker for positional reference.

To implant abdominal EMG electrodes, an incision 1 cm long was made in the skin below the ribcage to expose the oblique abdominal muscle. A small guide tube was then inserted into this incision and tunneled subcutaneously it reached the open scalp. Two coated stainless steel electrode wires (A-M Systems, #790500) were inserted through the tube until the ends were exposed though both incisions, allowing the tube to be removed while the wires remained embedded under the skin. Two gold header pins (Mill-Max Manufacturing Corporation, #0145-0-15-15-30-27-04-0) were adhered to the head bar with cyanoacrylate glue and the exposed wires between the header and neck incision were covered with silicone to prevent damage. Each wire exiting the abdominal incision was stripped of a section of coating and threaded through the muscle approximately 2 mm parallel from each other to allow for a bipolar abdominal EMG recording^46^. A biocompatible silicone adhesive (World Precision Instruments, KWIK-SIL) was used to cover the entry and exit of the muscle by the wires for implantation stability. The incision was then closed with a series of silk sutures (Fine Science Tools, #18020-50) and Vetbond (3M, #1469).

### Multiplane Imaging

To rapidly switch the focal plane between the brain and the skull, we integrated a ETL (Optotune, EL-16-40-TC-VIS-5D-C) into the laser path (SFig 2a). The ETL was placed adjacent to and parallel with the back aperture of the microscope objective (Nikon, CFI75 LWD 16X W) to maximize axial range, avoid vignetting^47^ and remove gravitational effects on the fluid-filled lens that could alter focal plane depth or cause image distortion^48^. An ETL controller (Gardasoft, TR-CL180) was used to control the liquid lens curvature. Pre-programmed steps in the curvature created rapid focal plane changes that were synchronized with image acquisition using transistor-to-transistor logic (TTL) pulses from the microscope. A microcontroller board (Arduino, Arduino Uno Rev3) was programmed to pass the first TTL pulse of every rapid stack to the ETL controller, which triggered a program that changed the lens curvature at predefined intervals (SFig 2b). The parameters of these steps were based on the framerate, axial depth, and number of images within the stack and were chosen to ensure the transitions of the lens’ curvature were done between the last raster scans of a frame and the beginning scans of the subsequent frame. The ability to trigger each rapid image stack independently using the microscope ensured consistent synchronization of the ETL and two-photon microscope even over long periods of data collection.

### Electrically tunable lens calibration

We calibrated the ETL-induced changes in focal plane against those induced by translation the objective along the Z axis (SFig 2). To generate a three-dimensional structure for calibration, strands of cotton were saturated with a solution of fluorescein isothiocyanate and placed in a 1.75 mm slide cavity (Carolina Biological Supply Company, #632255). These cotton fibers were then suspended in optical adhesive (Norland Products, NOA 133), covered with a glass cover slip, and cured with ultraviolet light (SFig 2d,e). At baseline, an ETL diopter input value of 0.23 was used as baseline as this generated a working distance closest to what would occur without an ETL. The objective was then physically stepped in the axial direction for 400 μm up and down in 5 μm steps, spanning 800 μm axially. The objective was then moved to the center of the stack and the diopter values were changed from −1.27 to 1.73 in 0.1 diopter steps while the objective was stationary, averaging 100 frames at each diopter value to obtain an image stack. The spatial cross-correlation between a single frame of the diopter stack and each frame of the objective movement stack were calculated to determine the change in focus location for each diopter value. This procedure was performed at three independent locations on the suspended fluorescein isothiocyanate cotton (SFig 2f). We performed calibrations of the magnitude across the usable range of ETL diopter values. While the difference in micrometers per pixel scaling relative to the baseline focal values was large across extremes in ETL-induced axial focal plane shift, the typical range used for imaging the brains of mice (<100 μm) had a negligible effect (approximately 0.01 µm/pixel) (SFig 2g).

To account for distortions within the focal plane, we imaged a fine mesh copper grid (SPI Supplies, 2145C-XA) (SFig 15). This square grid had 1000 lines per inch (19µm hole width, 6µm bar width). These values were used to determine the μm/pixel in the center of each hole in both the x and y direction. This allowed us to generate two three-dimensional plots of x, y, and μm/pixel points that were then fitted with a surface plot for distance calculations.

### EMG, locomotion, and respiration signals

EMG signals from oblique abdominal muscles were amplified and band pass-filtered between 300 Hz and 3 kHz (World Precision Instruments, SYS-DAM80). Thermocouple (Omega Engineering, #5SRTC-TT-K-20-36) signals were amplified and filtered between 2 and 40 Hz (Dagan Corporation, EX4-400 Quad Differential Amplifier) ^10^. The treadmill velocity was obtained from a rotary encoder (US Digital, #E5-720-118-NE-S-H-D-B). Analog signals were captured at 10 kHz (Sutter Instrument, MScan).

The analog signal collected from the rotary encoder on the ball treadmill was smoothed with a Gaussian window (MATLAB function: gausswin, σ = 0.98ms). EMG signal recorded from the oblique abdominal muscles from the mouse were filtered between 300 and 3000 Hz using a 5^th^-order Butterworth filter (MATLAB functions: butter, zp2sos, filtfilt) before squaring and smoothing (MATLAB function: gausswin, σ = 0.98ms) the signal to convert voltage to power.

The thermocouple signal was filtered between 2 and 40 Hz using a 5^th-^order Butterworth filter (MATLAB functions: butter, zp2sos, filtfilt) and smoothed with a Gaussian kernel (MATLAB function: gausswin, σ = 0.98ms).

### Abdominal pressure application

A custom-made pneumatically-inflatable belt (SFig 10a) was fabricated to directly apply pressure to the abdomen of mice. It consisted of three plastic bladders that were fully wrapped around the abdomen of mice. The belt was inflated with 7 psi of pressure to apply a steady squeeze for 2 seconds with 30 seconds of rest between squeezes to allow for a return to baseline (SFig 10b). The abdominal compression was oriented in such a way that no compression or tension was imparted to the spine longitudinally, as this could affect the results by pushing or pulling on the spine itself. Mice were observed with a behavioral video camera during imaging to check for potential compression-induced body positional changes and to monitor respiration.

### Motion tracking

Brain and fluorescent skull bead frames were deinterleaved. Each frame was then processed with a two-dimensional spatial median filter (3×3, MATLAB function: medfilt2). Occasionally, a spatial Gaussian filter (ImageJ function: Gaussian Blur) and contrast alterations (ImageJ function: Brightness/Contrast) were also applied prior to the median filter if the signal to noise ratio of the images resulted in poor tracking analysis.

At least three locations within the image sequence were chosen as targets for tracking. These template targets were manually selected regions of high spatial contrast (e.g. cell bodies) and were then averaged by pixel intensity across 100 frames during a period without brain motion to reduce noise for a robust matching template. Following the target template selection, a larger rectangular region of interest enclosing the template area was manually selected (MATLAB function: getpts) to spatially restrict the search (Fig 1d, SFig 4a).

For tracking, a MATLAB object was created (MATLAB object: vision.TemplateMatcher). A three-step search method was typically deployed at this step to increase computational speed for long image sequences. The sum of absolute differences between overlapping pixel intensities was calculated between the target and search windows, and the minimum value was chosen as the target position within the image. To monitor motion tracking, a displacement vector was then calculated that showed the motion in pixels between the current and prior image frames which was used to translate each image into a stabilized video sequence (MATLAB function: imtranslate). For visualization, a stabilized image was displayed alongside the target box displacement in the original image (MATLAB object: vision.VideoPlayer) to aid in manually checking for tracking failure.

Once the displacement in pixels was calculated for each target in a frame, the matrix of these values was searched for unique rows (MATLAB function: unique) to determine the number of unique target locations within the image. We then calculated the corresponding real distance between each unique location and the midlines of the image. A line was drawn between the image midline and the pixel location of the target. Then the calibration surface plot that depicts the calibration value in micrometers per pixel at each pixel for both x and y directions was integrated across this line (SFig11c, MATLAB function: trapz) to determine the distance in micrometers from the midline of the image. The real distance traveled between sequential frames was then calculated using these references by finding the difference of the target distances from the center of each frame. Performing the unique integrations first greatly increased the speed of processing the data. Motion was averaged across targets filtered with a Savitzky-Golay filter (MATLAB function: sgolayfilt) with an order of 3 and a frame length of 13 (Savitzky and Golay 1964). The standard error of the mean was calculated among the targets for each frame as well as the 90% probability intervals of the t-distribution (MATLAB function: tinv). The 90% confidence interval of the average object position in x and y was then calculated using the standard error of the mean and the probability intervals for the three signals at each frame (SFig 4b). The displacement of the fluorescent microspheres on the skull was then subtracted from the displacement of the brain to obtain a measurement of the motion of the brain relative to the skull.

### Motion direction quantification

We used principal component analysis (PCA) to find the primary direction of brain motion. Displacement data was first centered around the mean, then the covariance matrix of the positional data was calculated (MATLAB function: cov). The eigenvectors of this covariance matrix were then calculated (MATLAB function: eig) to determine the direction of the calculated principal components. To determine the magnitude of the vector, we took the mean of the largest 20% of the displacements from the origin (MATLAB function: maxk) (Fig 2a). This was done for each of the 316 recorded trials at 134 unique locations in 24 mice, where each trial is a continuous 10 minute recording. For locations with multiple trials, motion vectors were averaged to produce a single vector (Fig 2b, 5d, SFig 3b).

### MicroCT and vascular segmentation

A C57b/l6 mouse (male) was anesthetized with 5% isoflurane in oxygen and perfused a radiopaque compound (MICROFIL, MV-120) to label the vasculature. The mouse was then scanned with a microCT scanner (GE v|tome|x L300) at the PSU Center for Quantitative Imaging core from the nose to the base of the tail, covering 99.36 mm separated into 8280 slices with an isotropic pixel resolution of 12 μm. Images were collected using 75kV and 180 μA with aluminum filters for best contrast of tissue densities. Segmentation was done with 3D Slicer ^49^. Thresholding (3D Slicer function: thresholding) was first used to isolate the bone, and all voxels above a manually chosen intensity threshold were retained. Voxels that were preserved by the threshold tool but not required for the segmentation were removed within user-defined projected volumes (3D slicer function: scissors). The result was a high-resolution reconstruction of the skull, ribs, vertebrae, hips, and other small bones along the length of the mouse that retained their inner cavities. Segmentation of the vasculature surrounding the spine and skull was more difficult than isolating the bone because of the overlap in voxel intensity between the small vessels and the surrounding bone and tissues. The contrast agent also filled other organs (e.g. liver) with a similar intensity, so a simple threshold could not be used for the vasculature. We separated the vessels by using a freeform drawing tool (3D Slicer function: draw) to encapsulate the desired segmentation area for a single slice in two dimensions while ignoring unwanted similar contrast tissues. This process was repeated along the spine with a spacing of approximately 100 to 200 slices between labeled transverse areas. Once enough transverse freeform slices were created, they were used to create a volume by connecting the outer edges of consecutive drawn areas (3D Slicer function: fill between slices). This served as a mask that required all segmentation tools used to focus only on the voxels within the defined volume and ignore all others. The initial segmentation of the vasculature was created using a flood filling tool (3D Slicer function: flood filling). This tool labels vessels that are clearly connected within and across slices to quickly segment large branches of the network. The masking volume was utilized here to ignore connections to vessels or organs outside of the wanted space. The flood fill tool did not detect some connecting vessels, particularly ones located near the inner and outer surfaces of the vertebrae. In these instances, we utilized a segmentation tool that finds areas within a slice that shares the same pixel intensity around the entire edge (3D Slicer function: level tracing) to fill these gaps. In comparison to the bone, the three-dimensional reconstruction of the vessels was not smooth as they were smaller and had much more voxel intensity overlap with surrounding tissues and spaces. Thus, the segmentation was processed with a series of slight dilation operations that were followed by a matched erosion (3D Slicer function: margin). This technique of growing and shrinking the object repeatedly smoothed the surface and linked gaps between vessels. A specialized smoothing tool was then used for final polishing of the vasculature (3D Slicer function: smoothing).

### Brain motion simulations

Our calculations serve as a proof-of-concept. Thus, we selected an extremely simple geometric representation of the mouse CNS (Fig. 6). The brain and spinal cord (in pale pink) are surrounded by communicating fluid-filled spaces (in cyan). These consist of a central spherical ventricle internal to the brain and the subarachnoid space (SAS) on the outside of both brain and spinal cord. The SAS is connected to the ventricle by a straight central canal. In the center of the brain, above the ventricle, we placed a cavity meant to model the presence of the central sinus. In addition, we placed an outlet at the top of the skull to account for the fluid leakage out of the system through structures like the cribriform plate. The dimensions for system’s geometry in the reference (initial) state are reported in Supplementary Table 1. Like in Kedarasetti et al. (2022) ^15^, both brain and fluid-filled spaces are modeled as poroelastic domains: each consists of a deformable solid elastic skeleton through which fluid can flow. The two domains, which can exchange fluid, differ in the values of their constitutive parameters, the latter being discontinuous across the interface that separates said domains. All constitutive and model parameters adopted in our simulations are listed in Supplementary Table 1.

The governing equations have been obtained using mixture-theory ^15,50,51^ along with Hamilton’s principle ^52^, following the variational approach demonstrated in ^53^. Our formulation differs from that in ^53^ in that (i) each constituent herein is assumed to be incompressible in its pure form, and (ii) the test functions for the fluid velocity across the brain/SAS interface are those consistent with choosing independent pore pressure and fluid velocity fields over the brain and SAS, respectively. Hence, the overall pore pressure and fluid velocity fields can be discontinuous across the brain/SAS interface. The Hamilton’s principle approach allowed us to obtain consistent relations both in the brain and SAS interiors as well as across the brain-SAS interface. In addition, this approach yielded a corresponding weak formulation for the purpose of numerical solutions via the finite element method (FEM) (cf. ^54^).

By Ω_BR_ we denote the domain occupied by the cerebrum and spinal cord. By Ω_SAS_ we denote all fluid-filled domain, i.e., the SAS in a strict sense along with the central canal and the ventricle. These domains are time dependent. We denote the interface between Ω_BR_ and Ω_SAS_ by Γ. The unit vector *m* is taken to be normal to Γ pointing from Ω_BR_ into Ω_SAS_. Subscripts s and f denote quantities for the solid and fluid phases, respectively. In their pure forms, each phase is assumed incompressible with constant mass densities 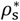 and 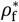.Then, denoting the volume fractions by *ϕ*_s_ and *ϕ*_f_, for which we enforce the saturation condition *ϕ*_s_ + *ϕ*_f_ = 1, the mass densities of the phases in the mixture are 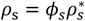 and 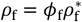.The symbols *u*, ***ν***, and T (each with the appropriate subscript), denote the displacement, velocity, and Cauchy stress fields, respectively. The quantity ***ν***_flt_ = *ϕ*_f_(***ν***_f_ − ***ν***_s_) is the filtration velocity. The pore pressure, denoted by *p*, serves as a multiplier enforcing the balance of mass under the constraint that each pure phase is incompressible. To enforce the jump condition of the balance of mass across Γ, we introduce a second multiplier, denoted ℘. The notation ⟦*a*⟧ indicates the jump of *a* across Γ. We choose the solid’s displacement field so that ⟦*u*_s_⟧ = 6 (i.e., *u*_s_ is globally continuous). Formally, ***ν***_f_ and *p* need not be continuous across Γ. Possible discontinuities in these fields have been the subject of extensive study in the literature (cf., e.g., ^53,55,56^) and there are various models to control their behavior (e.g., often ***ν***_flt_ and *p*_f_ are constrained to be continuous ^56^). We select discontinuous functional spaces for *p* and ***ν***_f_ and we control their behavior by building an interface dissipation term in the Rayleigh pseudo-potential in our application of Hamilton’s principle (similarly to ^53^). This dissipation can be interpreted as a penalty term for the discontinuity of the filtration velocity. Before presenting the governing equations, we introduce the following two quantities: *k*_f_ = (1/2)*ρ*_f_***ν***_f_ *·* ***ν***_f_ (kinetic energy of the fluid per unit volume of the current configuration) and *d* = *ρ*_f_(***ν***_f_ − ***ν***_s_) *· m*, which the jump condition of the balance of mass requires to be continuous across Γ.

The strong form of the governing equations, expressed in the system’s current configuration (Eulerian or spatial form; cf. ^57^) are as follows:

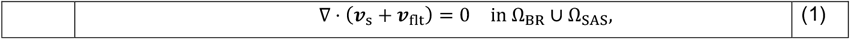

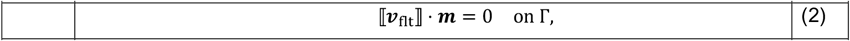

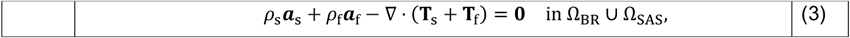

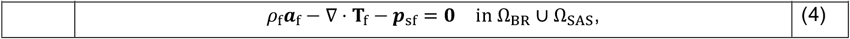

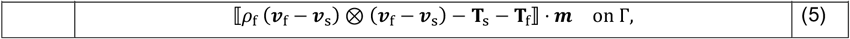

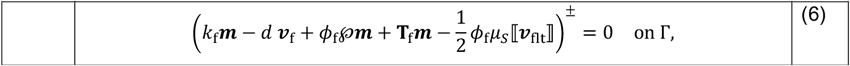

where *a*_s_ and *a*_f_ are material accelerations, the superscript ± refers to limits approaching each side of the interface, *μ*_*s*_ is a viscosity like parameter (with dimensions of velocity per unit volume) characterizing the dissipative nature of the interface, and where the terms **T**_s_, **T**_f_, and ***p***_sf_ are governed by the following constitutive relations

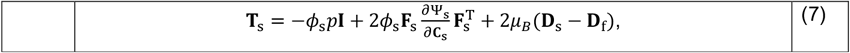

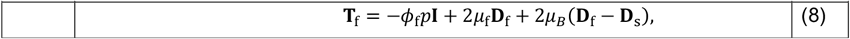

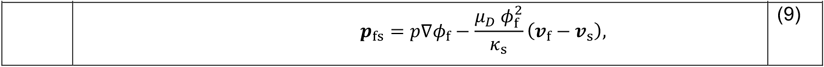

where Ψ_s_ is the strain energy of the solid phase per unit volume of its reference configuration, **F**_s_ = **I** + ∇_s_***u***_s_ is the deformation gradient with ∇_s_ denoting the gradient relative to position in the solid’s reference configuration, 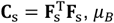 is the Brinkmann dynamic viscosity, **D** = (∇***ν***_s_)_sym_, **D**_f_ = (∇***ν***_f_)_sym_, (∇***ν***)_sym_ denoting the symmetric part of ∇***ν***, *μ*_f_ is the traditional dynamic viscosity of the fluid phase, *μ*_*D*_ is the Darcy viscosity, and *κ*_s_ is the solid’s permeability. For Ψ_s_ we choose a simple isochoric neo-Hookean model: 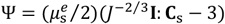,where *J* = det **F**_s_ and 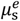 is the elastic shear modulus of the pure solid’s phase. It is understood that the constitutive parameters in Ω_BR_ are different from those in Ω_SAS_.

The details of the boundary conditions and of the finite element formulation are provided in the supplementary materials. Here we limit ourselves to state that the problem is solved by using the motion of the solid as the underlying map of an otherwise Lagrangian-Eulerian formulation for which the reference configuration of the solid phase serves as the computational domain. The loading imposed on the system consists of a displacement over a portion of the dural sac of the spinal cord we denote as SZ (for the squeeze zone), meant to simulate a squeezing pulse provided by the VVP. This displacement is controlled so that a prescribed nominal uniform squeezing pressure is applied to the said zone. Flow resistance boundary conditions are enforced at the outlet at the top of the skull, and a resistance to deformation is also imposed on the walls of the central sinus.

**Supplementary Table 1.**
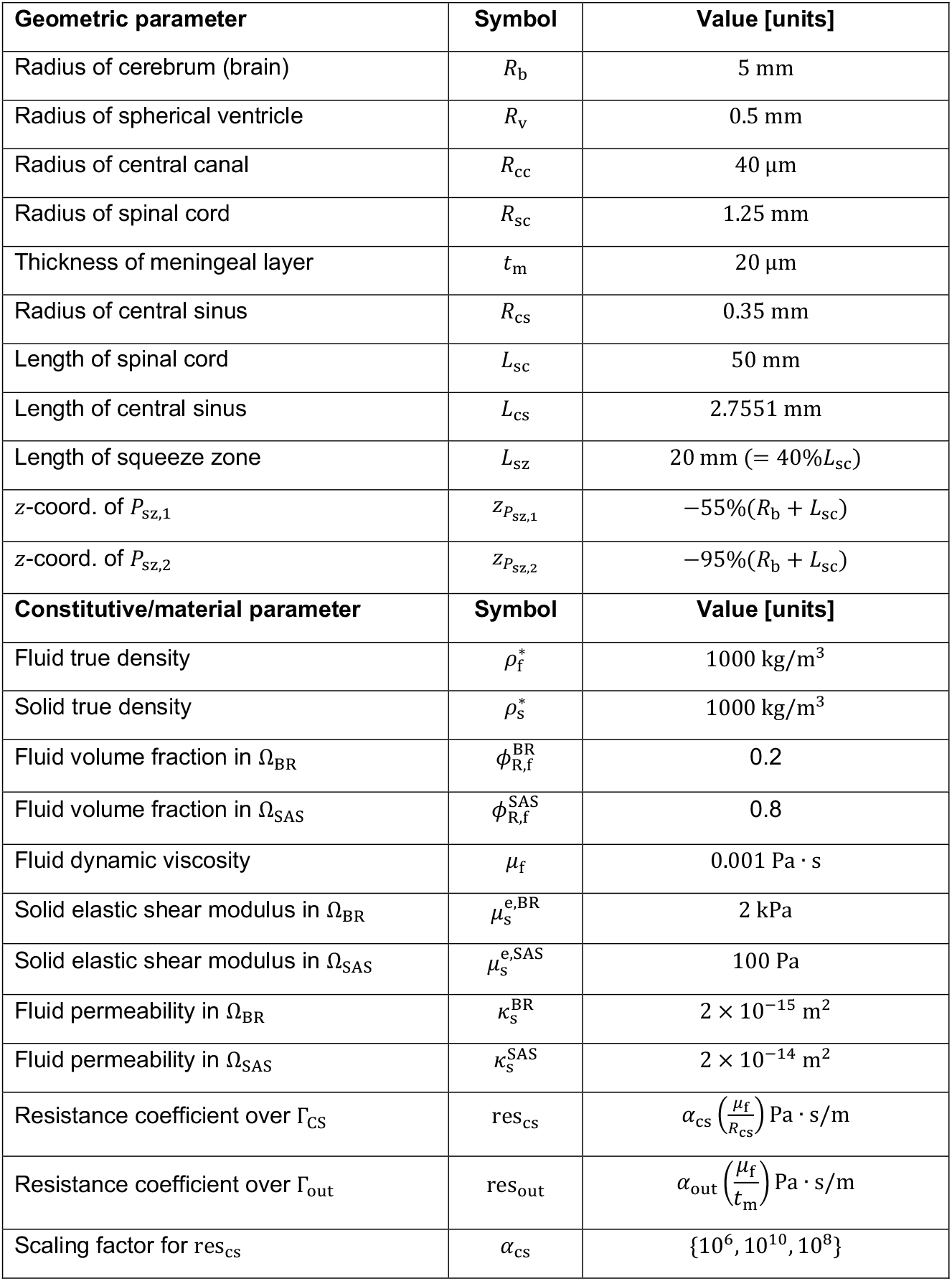
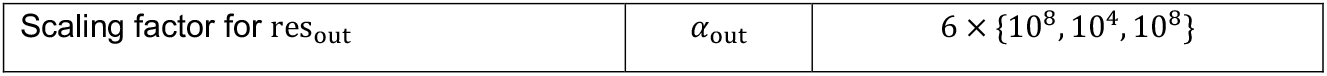
Simulation geometry data and constitutive parameters. Dimensions of the central sinus compartment adopted in this geometry (i.e., *R*_cs_ and *L*_cs_) were chosen to match the reference volume of a cylindrical central sinus with radius and length of 1*5*0 μm and 1*5* mm, respectively.

### Supplementary Material — Finite Element Formulation and Boundary Conditions Notation

Here we report the weak form of the governing equations. To avoid proliferation of symbols, given a field ζ, the test function for that field will be denoted by 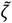.We denote the reference configuration of the solid phase by Ω_s_.We formulate the weak form of our problem over Ω_s_. Let Ω(*t*) = Ω_BR_ ∪ Ω_SAS_ denote the current configuration of the (entire) system. We denote by ***X***_s_ points in Ω_s_. Denoting the motion of the solid by ***χ***_s_(***X***_s_, *t*), under common assumptions from mixture theory, ***χ***_s_ is a smooth map with smooth inverse from Ω_*s*_ to Ω(*t*). The gradients over Ω(*t*) and Ω_s_will be denoted by ∇ and ∇_s_, respectively. Given a quantity ζ(***x***, *t*) over Ω(*t*), ζ^*σ*^ is defined as ζ^*σ*^(***X***_s_, *t*) = ζ(***χ***_s_(***X***_s_, *t*), *t*). The fields *u*_s_, **F**_s_, and *J*_s_are understood to have Ω_s_ as their domain. Given any two fields ζ and *φ* over some domain Θ such that their (pointwise) inner-product is meaningful, we denote by (ζ, *φ*)_Θ_ the integral over Θ of said inner product. We denote by Γ_s_, the inverse image of the brain-SAS interface under the solid phase motion. The notation 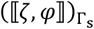 will indicate the integral over Γ_s_ of the jump of the inner-product of ζ and *φ* across Γ_s_.

### Weak Form

For ease of writing, the weak form shown here is written assuming that *u*_s_ and 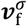 are prescribed on the external boundary of the system. The boundary conditions are indicated in the following subsection. The weak form is as follows:

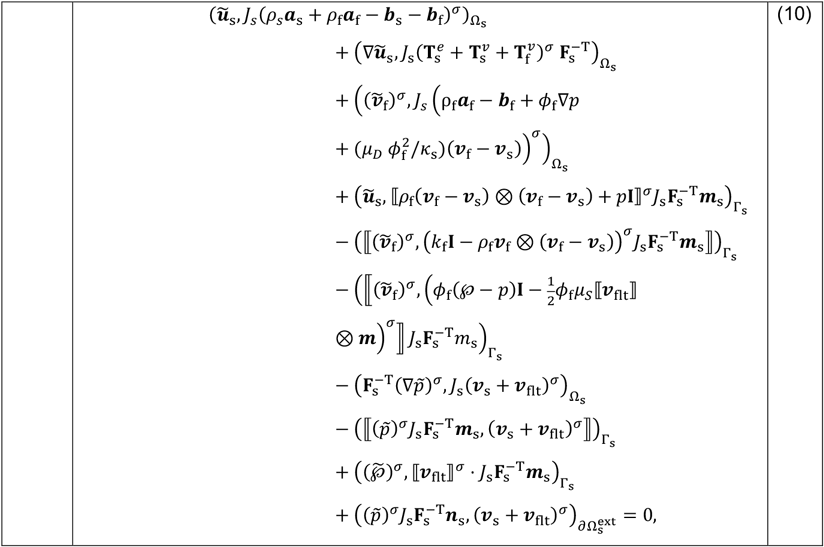

where 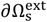 denotes the outer-most boundary of Ω and ***n***_s_ is the associated outward unit normal. The above weak form, modified to enforce the boundary conditions listed later, is required to hold for all test functions 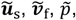 and 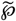 in functional spaces chosen in a coordinated manner to the functional spaces selected for the unknown fields *u*_s_, ***ν***_f_, *p*, and ℘. As a formal analysis concerning the well-posedness of the problem considered in this paper has yet to be developed, we avoid characterizing the spaces in question using the formal language of Sobolev spaces. Rather, we limit ourselves to describing the details of our practical implementation. With the few exceptions that we will describe next, our implementation follows standard practices in the FEM literature on solid and fluid mechanics (cf. ^54^).

As mentioned in the main body of the paper, *u*_s_ is globally continuous over Ω_s_. Its numerical representation was done using a second-order Lagrange polynomial FE field. The fields ***ν***_f_ and *p* were taken to be continuous over the subsets of Ω_s_ corresponding to the brain and the SAS. However, these fields are not continuous across Γ_s_. The FE fields taken to interpolate ***ν***_f_ and *p* were second-order and first-order Lagrange polynomials, respectively. The field ℘ was taken to be continuous over Γ_s_ (this field does not exist away from Γ_s_) and interpolated using first-order Lagrange polynomials.

### Note on Integration by Parts

The weak enforcement of Eq. (1), namely the continuity equation for this problem, was done by testing said equation by 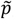,integrating the resulting form over the problem’s domain, and applying integration by parts. This treatment of the continuity equation is not standard. The rationale for this approach is the desire to avoid approximating the gradient of the volume fraction *ϕ*_*f*_. This choice has additional consequences in the treatment of the momentum equations and any boundary condition involving boundary tractions. In the momentum equations, we do *not* apply integration by parts to terms involving the gradient of the pore pressure. When it comes to boundary tractions, as it would be physically incorrect to prescribe pore pressure boundary values, we retain the associated pore pressure in the boundary contributions.

### Note on Implementation of Boundary Conditions Involving Traction

Here we indicate boundary conditions involving tractions in the *current configuration* of the system. This is done to facilitate the readability. As indicated earlier, the motion of the solid phase provides the ALE map needed for the pullback of said conditions to the actual computational domain. This said, we note that our computations were carried out using COMSOL Multiphysics® (v. 6.1. www.comsol.com. COMSOL AB, Stockholm, Sweden). The latter provides automatic support for these operations. That is, a user can specify whether a contribution to a weak form is to be evaluated in the “Spatial” frame (here Ω(*t*)) or the “Material” frame (here Ω_s_). We have taken advantage of this feature in our calculations.

### Boundary Conditions

With reference to Fig. 6, the overall geometry of the system is axially symmetric and a cylindrical coordinate system is defined such that the *z* axis is the dashed line in the figure with the positive direction from the tail towards the head. The radial coordinate *r* is in the direction perpendicular to the *z* axis. The boundary of the brain-SAS over which boundary conditions are applied consists of the surface Γ_CS_ surrounding the central sinus, and of the union of the subsets Γ_out_,Γ_ext_, and Γ_SZ_. Γ_out_ is an outlet /inlet meant to represent a structure like the cribriform plate through which CSF can exit/enter the system. Γ_SZ_ is the region on which the squeezing action of the VVP onto the dural sack is applied. Γ_ext_ denotes the remaining portion of the SAS external boundary. Axial symmetry was enforced in a standard fashion, namely requiring the radial component of vector fields *z*-axis. The rest of the boundary conditions are as follows:

- *u*_s_ = 0 on Γ_ext_ ∪ Γ_out._
- *u*_s_ = −*u*_0,rad_(*t*)*f*_SZ,space_(*z*)*e*_*r*_ on Γ_SZ_, where
  - *f*_SZ,space_(*z*) is a (unit) step function over the spatial interval 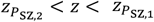 smoothed so to be continuous up to 2^nd^ order derivatives over transition zones 10% in size of the function’s support.
  - *u*_0,rad_ is a positive scalar function of time subject to the following constraint: 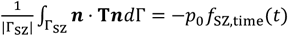,where T is the total Cauchy stress acting on the mixture (i.e., solid and fluid phases combined), ***n*** is the outward unit normal in the current configuration on Γ_SZ_, *p*_0_ a prescribed pressure value, and *f*_SZ,**t**ime_(*t*) a unit step function over the time interval 0 < *t* < *t*_squeeze_, smoothed so to be continuous up to 2^nd^ order derivatives over a transition zones 10% in size of the function’s support. That is, *u*_0,rad_ (*t*) was controlled so that the spatial average of the normal traction over the SZ was equivalent to a uniform pressure distribution of value *p*_0_.
- ***ν***_f_ = ***ν***_s_ on Γ_ex**t**_ ∪ Γ_SZ_ ∪ Γ_CS_ — This is a “no slip” boundary condition for the fluid relative to solid phase. This boundary condition has been enforced weakly (cf., e.g., ^59^).
- Robin boundary condition on Γ_CS_ — This boundary condition is meant to allow the central sinus (CS) to deform in response to intracranial pressure changes as well as brain movement. Physiologically, this response is mediated by blood flow in the CS. We have modeled this response through a traction distribution on Γ_CS_ proportional to the velocity of Γ_CS_: T***n*** = −res_cs_ ***ν***_s_, where, again, T is the total Cauchy stress on the mixture, ***n*** is the outward unit normal, and where ζes_cs_ is a resistance constant indicated in Table 1. We have investigated the effects of a range of values of this constant.
- Robin boundary condition on Γ_out_: This is a boundary condition meant to model the outflow of CSF from the skull through pathways like the cribriform plate and the olfactory nerves. In our simulations we have not included sources of production of CSF. Hence, the condition on Γ_out_ is bidirectional, i.e., it allows for both outflow and inflow of CSF. This condition amounts to a hydraulic resistance, which we have implemented as a Robin boundary condition. Specifically, we have enforced the following condition on Γ_out_: T_f_***n*** = −ζes_out_ ***ν***_flt_, where T_f_ is the total Cauchy stress on the fluid phase, ***n*** is outward unit normal, and ζes_out_ is a constant hydraulic resistance indicated in Table 1. As for ζes_cs_, we have investigated the effects of different values of this constant.

### Note on Computer Implementation

The mesh and solver were developed using the standard facilities available in COMSOLMultiphysics®. We have employed a mesh consisting of 63180 triangles and 53792 quadrilaterals for a total of 116972 elements. Eight boundary layers with a stretching factor of 1.2 have been placed along the brain-SAS interface. The total number of degrees of freedom is 1,487,327: 690014 for *u*_s_, 446450 and 257014 for ***ν***_f_ in the brain and SAS, respectively, 56669 and 33816 for *p* in the brain and SAS, respectively, 3363 for ℘. Finally there is one degree of freedom for *u*_o,rad_. Time integration was carried out using a variable step/variable order BDF ^60^ method, with order raging from 2 to 5 and with a maximum time step set to 0.001s. The maximum time used for the computations was 10s, to simulate the 2s −squeeze pulse along with the recovery phase of the system after the squeeze ends. The solver was fully coupled and monolithic. MUMPS was selected as the algebraic solver.

**Supplementary Figure 1.**
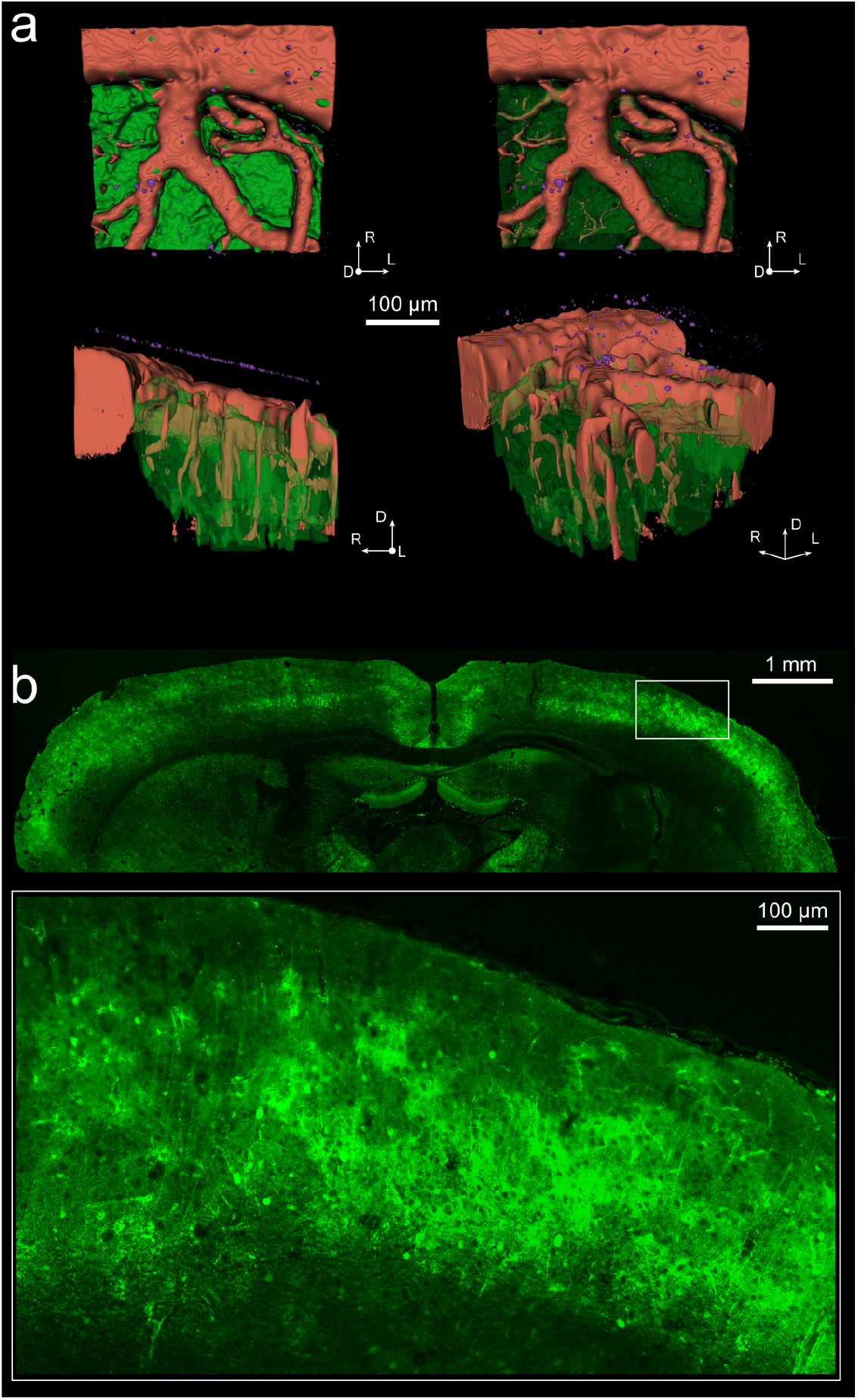
Microspheres and brain. **a**. Reconstruction of GFP-expressing parenchyma (green), blood vessels (red), and fluorescent microspheres (magenta). The axes are labeled dorsal (D), rostral (R), and lateral (L). Penetrating vessels can be seen through the semi-transparent brain in the bottom left and bottom right images. **b**. Coronal section of a GFP-expressing mouse brain, showing ubiquitous labeling of cells.

**Supplementary Figure 2.**
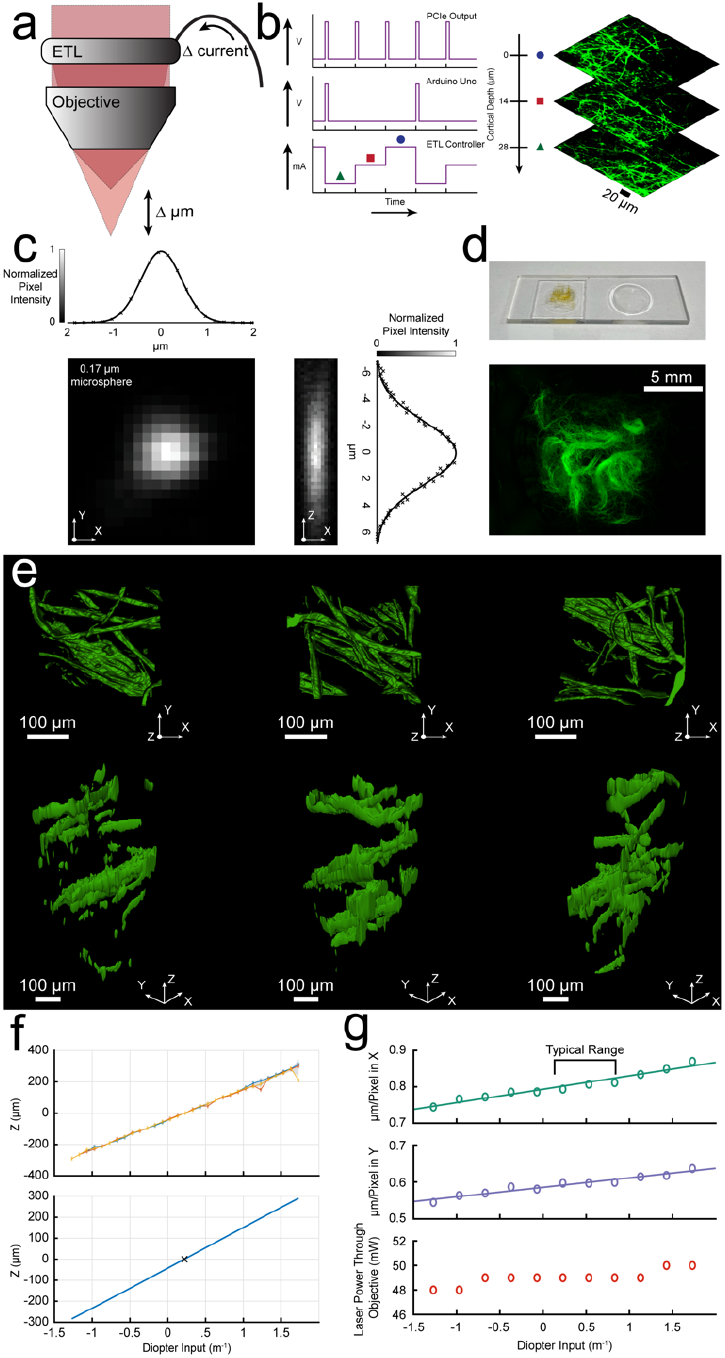
Axial calibration of electrically-tunable lens. **a**. A change in the current input to the lens generates a curvature change in the lens, which alters the focus. **b**. Synchronization of ETL focus change with microscope scanning. A TTL pulse is generated at the beginning of each frame from the PCIe board in the computer controlling the microscope (top left). An Arduino Uno was programmed to filter all pulses besides the first of the stack (middle left). This pulse was then sent to the ETL controller to prompt a predetermined set of current steps that were sent to the ETL (bottom left). These currents changes created a rapid stack with each depth captured as a single frame (right). **c**. The point spread function in the X (left) and Z (right) directions of the two-photon microscope created with a 0.17µm fluorescent microsphere and a 0.8 NA N16XLWD-PF 16x Nikon objective. The ETL obscures part of the back aperture, resulting in a lower effective NA. **d**. Calibration of the ETL focal range. To provide a fluorescent three-dimensional structure, cotton stands were dipped in a solution of fluorescein isothiocyanate and suspended in optical adhesive within a concave slide. **e**. Three-dimensional segmentations created using fluorescent cotton strands from three locations (left to right). **f**. Calibration of the ETL diopter shifts to focal plane shifts. Three locations in the cotton (shown in **e**) were imaged by shifting the ETL focus and by translating the object in Z and aligned by correlational matching of images (top). These averages are potted for each location in colored lines with the shaded standard deviation. The linear regression is also plotted as a solid blue line, with zero µm being the focus neutral diopter value (bottom). **g**. From top to bottom, change in X and Y scaling and laser power as a function of diopter value. Changing the diopter of the ETL had negligible changes in magnification and laser power output in the typical imaging range.

**Supplementary Figure 3.**
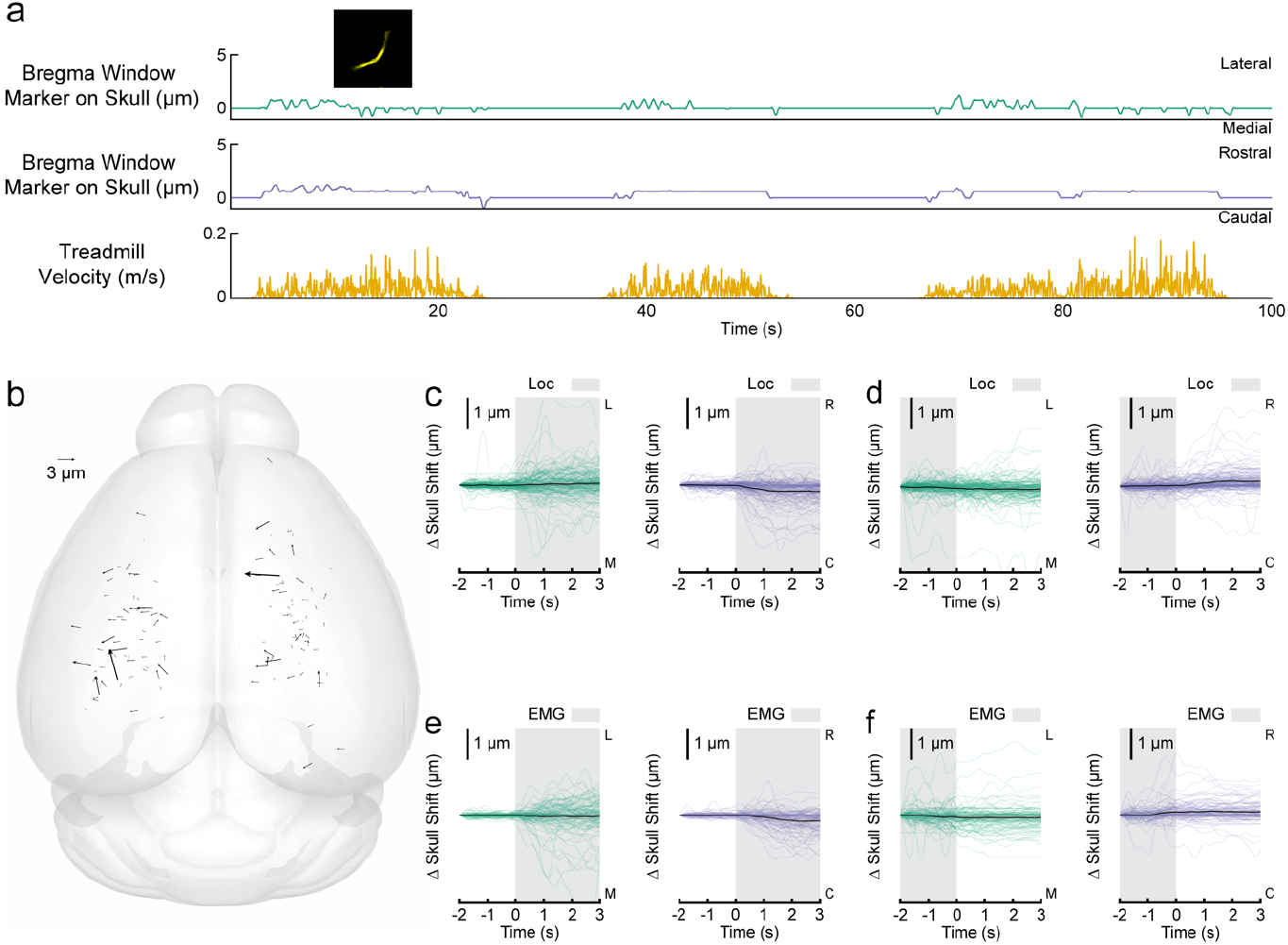
Negligible skull motion during locomotion. **a**. ‘Worst-case’ skull motion in a 55 gram mouse. A fluorescent marker on the skull at bregma was imaged due to its large distance from the implanted head bar (implanted caudally of lambda) to maximize the ability for the skull to displace during locomotion. **b**. A plot of skull displacement, calculated from the same trials as the brain motion (N=134 sites in 24 mice). Note the small size and lack of clear direction. **c**. Locomotion-triggered average skull motion for each trial. The black line shows the mean, and the shaded portion denotes 90 percent confidence interval. **d**. Locomotion cessation-triggered average skull motion. **e**. EMG-triggered average skull motion **f**. EMG cessation-triggered average skull motion.

**Supplementary Figure 4.**
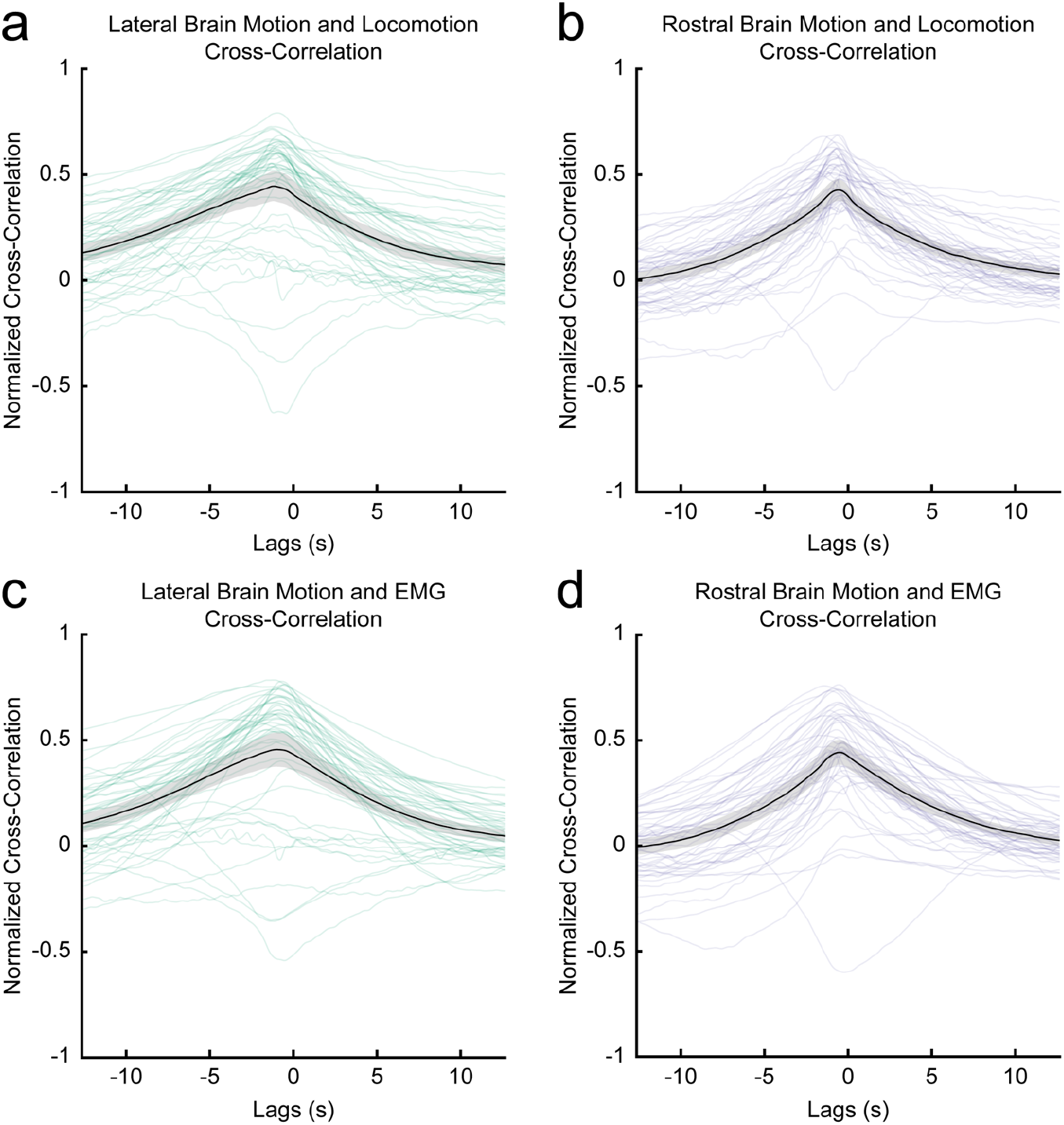
Cross-correlations between cortical brain motion, locomotion and abdominal EMG. **a**. Cross-correlation between locomotion and lateral cortical motion. Black line shows mean, with shading showing 90 percent confidence interval. **b**. Cross-correlation between locomotion and rostral cortical motion. **c**. Cross-correlation between EMG and lateral cortical motion. **d**. Cross-correlation between EMG and rostral cortical motion.

**Supplementary Figure 5.**
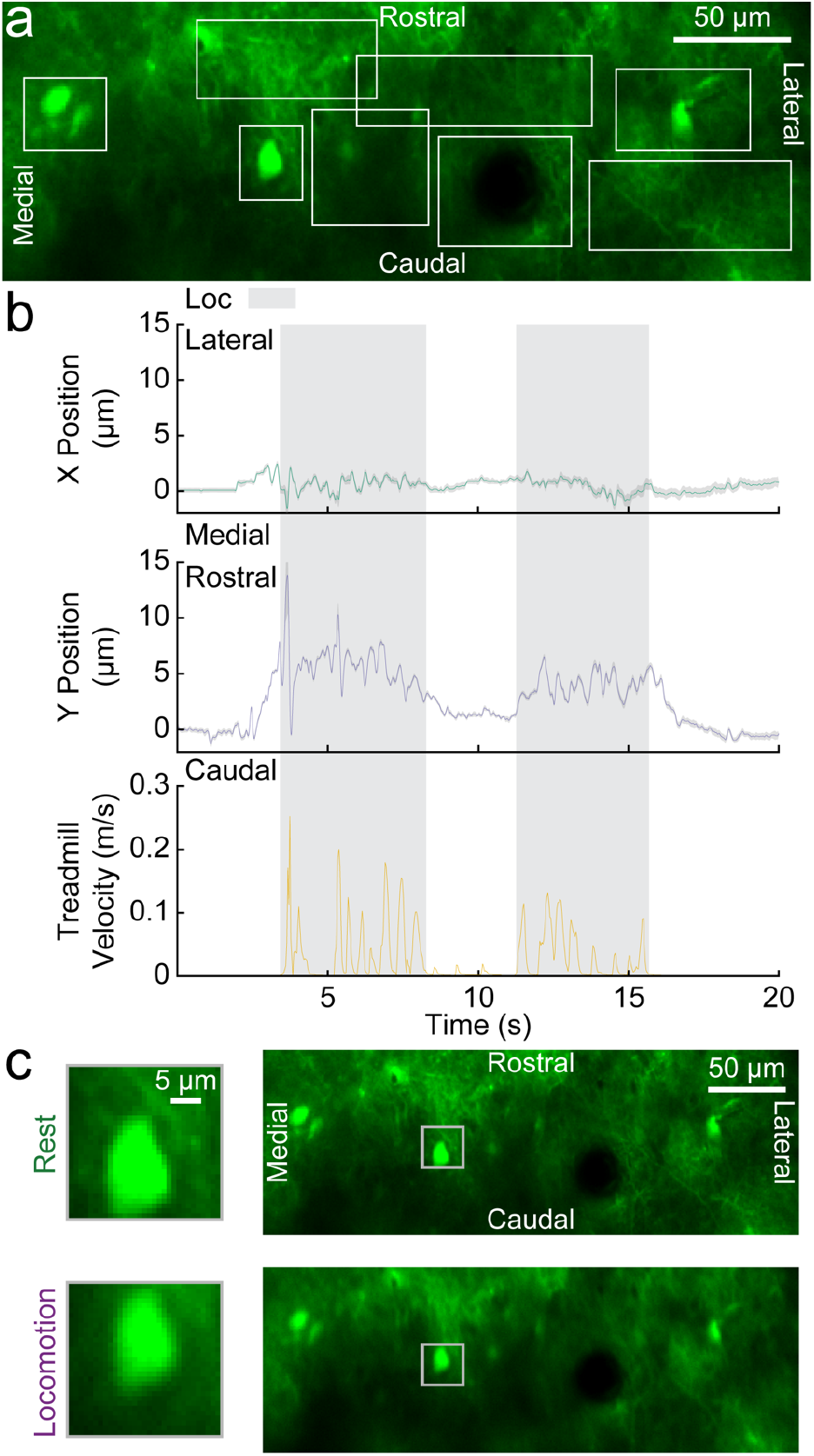
Template-matching algorithm used to track the brain is robust across the field of view. **a**. An image of the GFP-expressing parenchyma. Each of the eight bounding boxes (white) represents a tracking template area for the matching algorithm to follow. **b**. The targets were tracked at each of the eight locations and the mean and 90 percent confidence interval (shading) were calculated and plotted. The tight confidence interval bounds highlight the confidence in tracking different structures at various locations within the image as well as a lack of brain distortion within the field of view, indicating ridged translation. **c**. Images of the brain (from **a**) when the mouse is at rest (top) and during a locomotion event (bottom). The neuron seen in the bounding box (gray) displaces rostrally and laterally during locomotion when compared to its resting position.

**Supplementary Figure 6.**
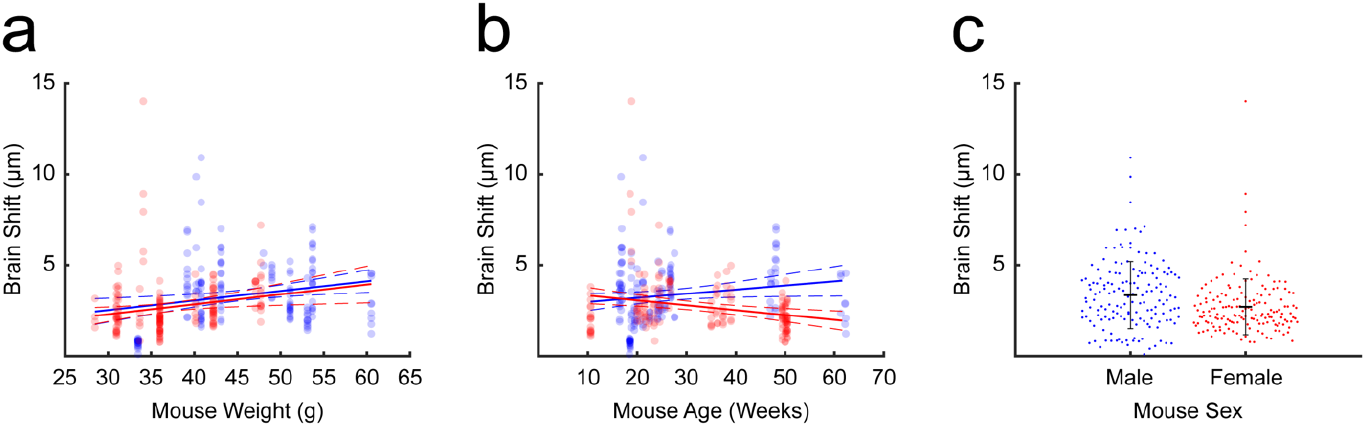
The impacts of sex, age, and weight on measured brain motion. **a**. Magnitude of brain displacement within the skull plotted as a function of mouse weight. The solid blue line represents the linear fit for males (p = 0.0064, R^2^ = 0.0478) and the solid red line represents the linear fit for females (p = 0.0145, R^2^ = 0.0368). Dashed lines show the 95 percent confidence intervals. **b**. Magnitude of brain displacement within the skull as a function of mouse age. The solid blue line represents the linear fit for males (p = 0.0502, R^2^ = 0.0250) and the solid red line represents the linear fit for females (p = 0.0015, R^2^ = 0.0609). Dashed lines show the 95 percent confidence intervals. **c**. Brain displacement for males and females. Bars show the mean and the standard deviation. A two-sample Kolmogorov-Smirnov test on these data sets rejects the null hypothesis that these sets are from the same continuous distribution at a 5 percent significance level (p = 0.00002).

**Supplementary Figure 7.**
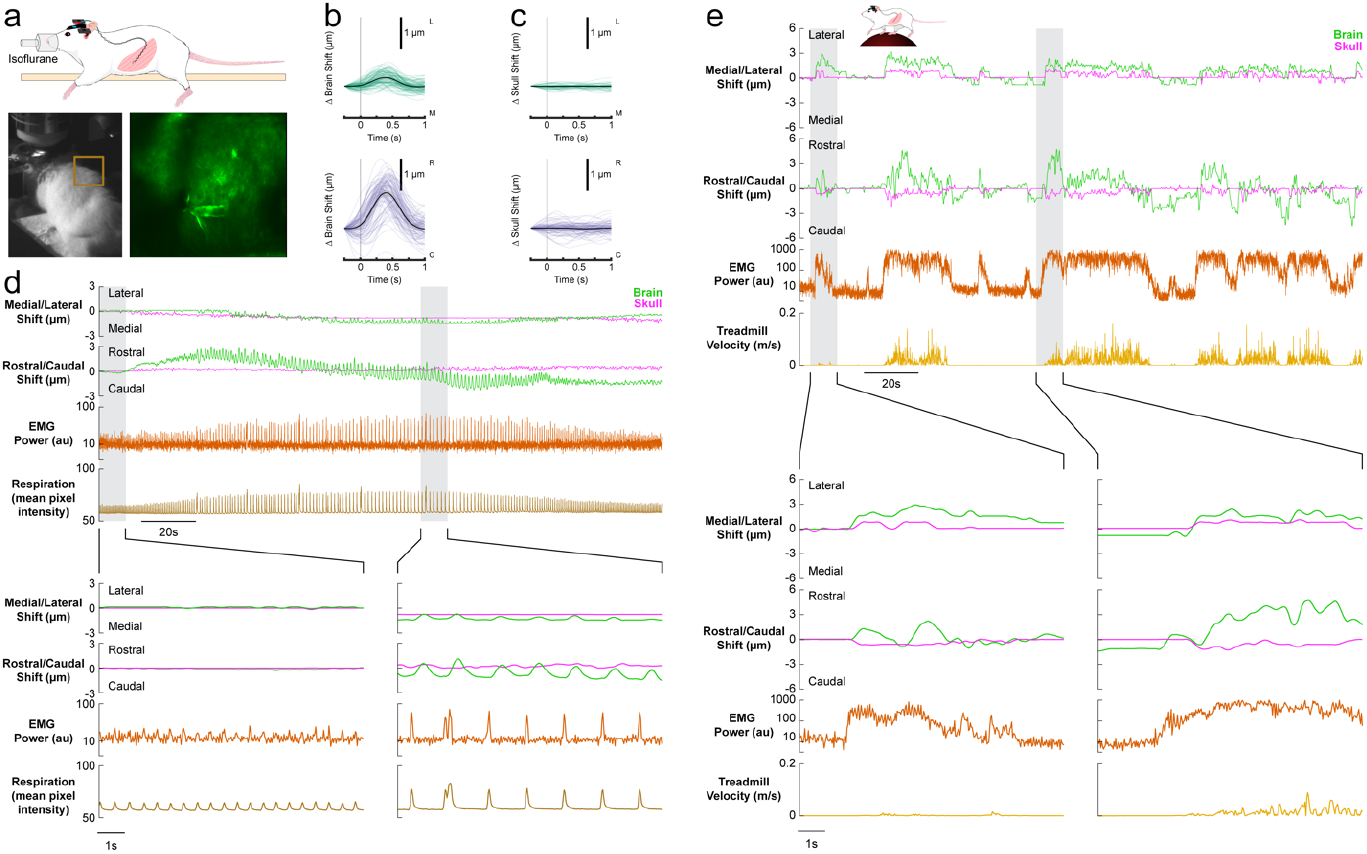
Respiration-driven brain motion is only observed under deep anesthesia in mice when abdominal muscles are engaged. **a**. The respiration of the mouse anesthetized with isoflurane and instrumented with abdominal EMG electrode was monitored using a behavioral camera by measuring the mean pixel intensity of a box drawn across the edge of the body (box in bottom left image). Inset shows brain visualized under the two-photon microscope (bottom right). **b**. EMG-triggered motion during period of deep anesthesia respiration trial in the medial/lateral (top) and rostral-caudal (bottom) directions. The black line shows the mean with a shaded 90 percent confidence interval. **c**. EMG-triggered skull motion **d**. Brain (green) and skull (magenta), abdominal EMG power (orange) and behavioral camera respiration (brown) during varying levels of anesthesia. Isoflurane concentration began at 0.5% at 5 seconds, then was increased to 5% in oxygen for 120 seconds, after which it was reduced to 0.5% in oxygen once again. Light anesthesia is characterized by shallow breaths with minimal abdominal muscle contraction and produced no detectable pattern of brain motion within the skull (lower left). Deep anesthesia, characterized by slower and deeper breaths, resulted in increased abdominal muscle activation and brain motion within the skull (lower right). **e**. The same location was imaged again in the same mouse on a subsequent day. Locomotion drove larger brain (green) and skull (magenta) displacements compared to respiration-induced brain motion under anesthesia (shown in **d**). The abdominal muscle power (orange) also shows much stronger abdominal muscle contractions during locomotion events (gold). Abdominal muscle activation without a locomotion event (lower left) still resulted in a rostro-lateral brain displacement within the skull.

**Supplementary Figure 8.**
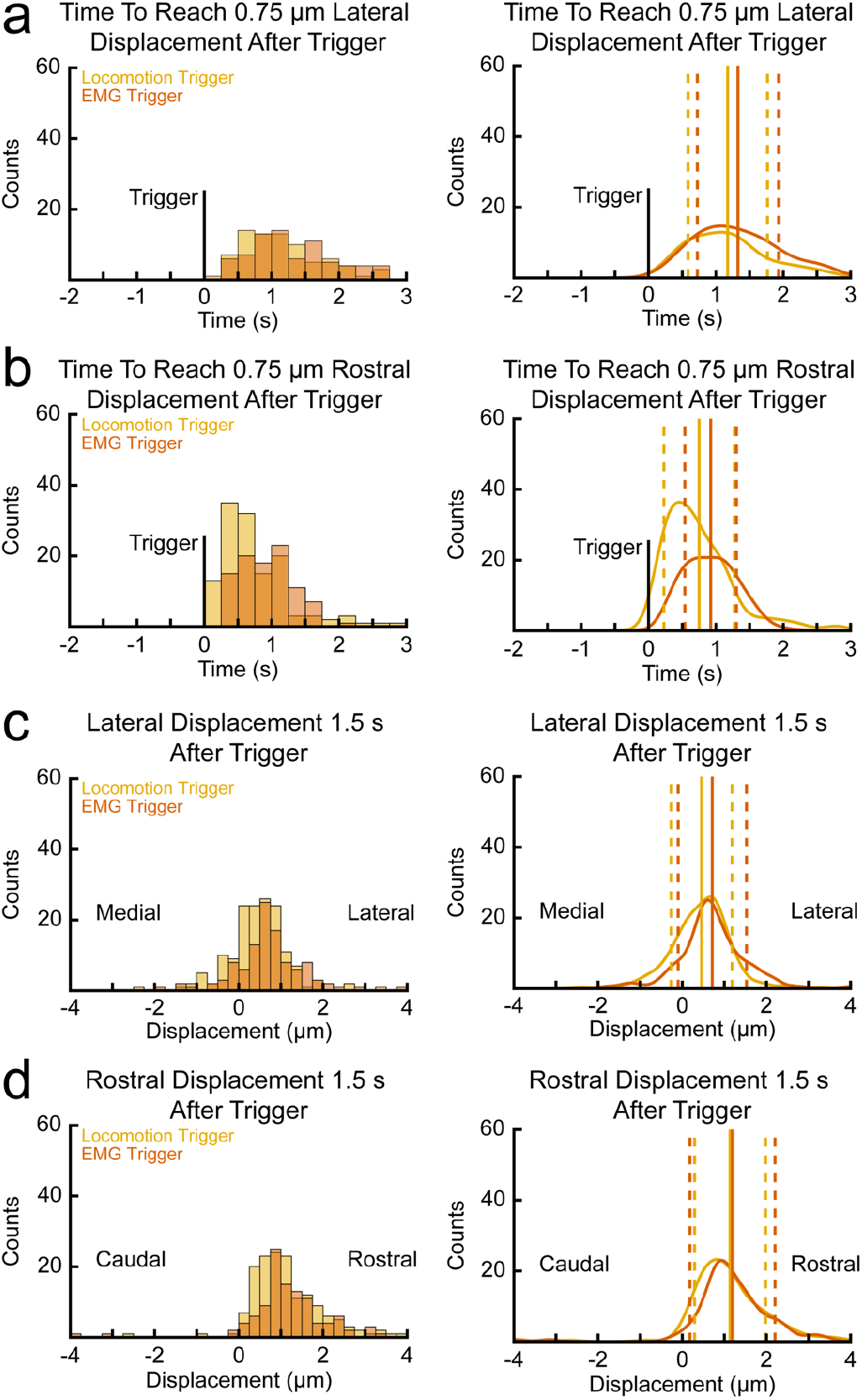
The brain displaces more quickly following an electromyography event than a locomotion event. **a**. The time that the brain takes to displace laterally 0.75µm following a locomotion and EMG event onset as histograms (left) and as probability density functions with corresponding means and standard deviations (right). **b**. The time that the brain takes to displace rostrally 0.75µm following a locomotion and EMG event onset as histograms (left) and as probability density functions with corresponding means and standard deviations (right). **c**. The distance the brain has displaced laterally 1.5 seconds after a locomotion and EMG event onset as histograms (left) and as probability density functions with corresponding means and standard deviations (right). **d**. The distance the brain has displaced rostrally 1.5 seconds after a locomotion (black) and EMG (orange) event onset as histograms (left) and as probability density functions with corresponding means and standard deviations (right).

**Supplementary Figure 9.**
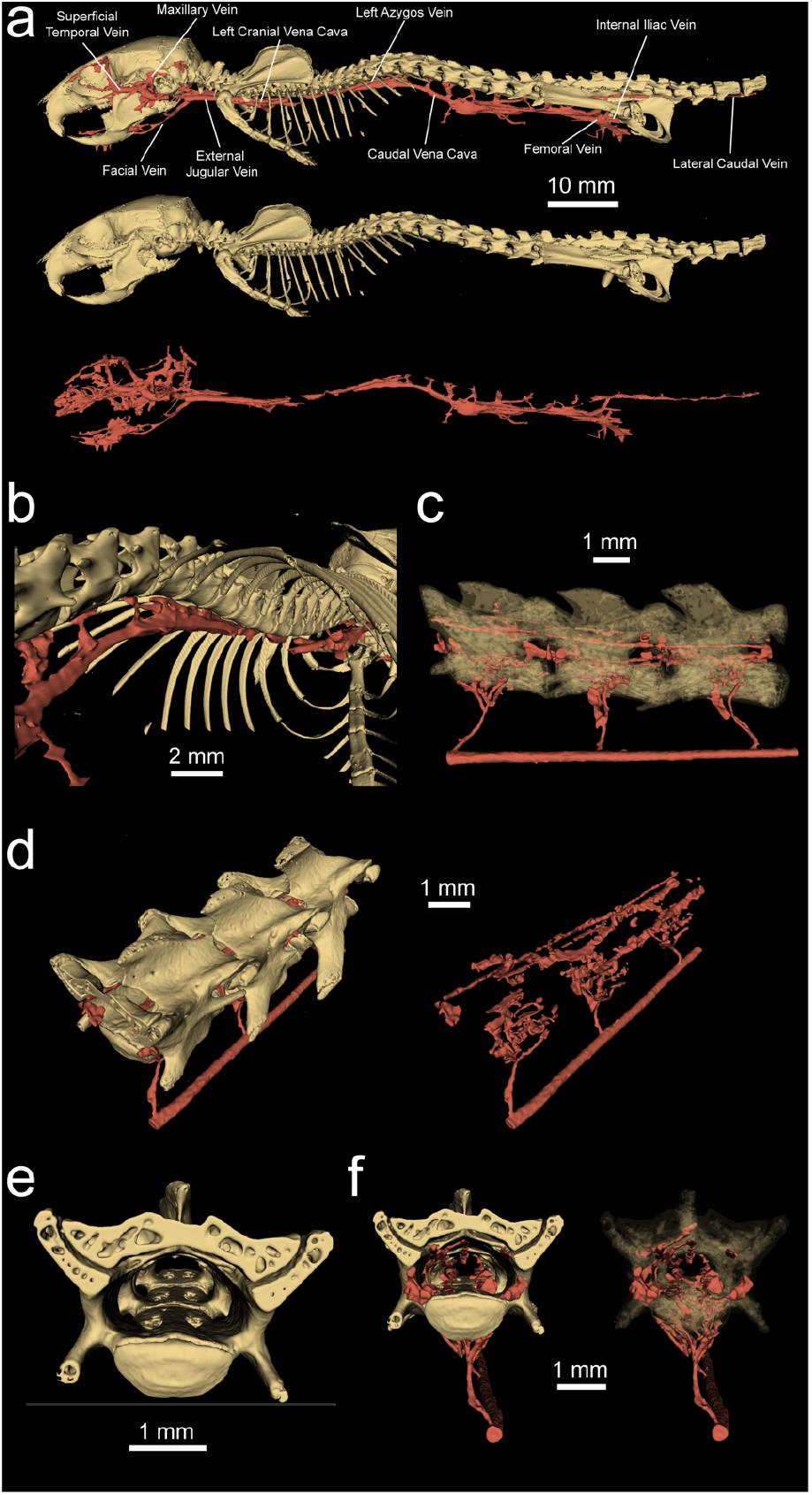
MicroCT imaging of spine and associated vasculature. **a**. The skeleton and spinal vasculature. **b**. A view of the vessels within the rib cage. The gap observed between the caudal vena cava and both cranial vena cava is occupied by the heart, which was not included. Note the lack of connections between the caudal vena cava and the vertebrae within the rib cage. **c**. A view of the L3, L4, and L5 vertebrae showing connections between the caudal vena cava and VVP within the vertebrae. **d**. The internal VVP is shown both with bone (left) and without bone (right). **e**. Two holes are present on the internal ventral surface of the vertebrae. These may act as pathways for veins in the abdomen to connect to the VVP within the lumbar section of the vertebral column. **f**. Visualization of the veins and vertebrae with a focus on the internal ventral holes in the bone. Veins occupy the holes in the vertebrae, which can be seen both when the bone is opaque (left) and semi-transparent (right).

**Supplementary Figure 10.**
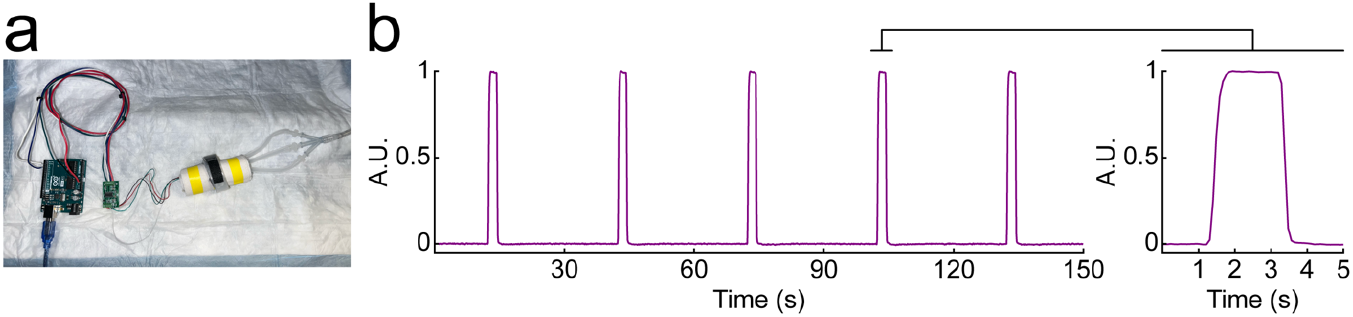
Repeatable pressures applied by the inflatable belt. **a**. An Arduino Uno collected data from a strain gauge wrapped with paper towels and inserted into the inflatable belt used for abdominal compressions. **b**. The belt was inflated with 7 psi of compressed air for two seconds at 30 second intervals. The resulting pressure applied to the strain gauge was consistent in intensity and duration (left). A closer look at a single compression demonstrates the rapid onset and offset transients of the pressure applied (right).

**Supplementary Figure 11.**
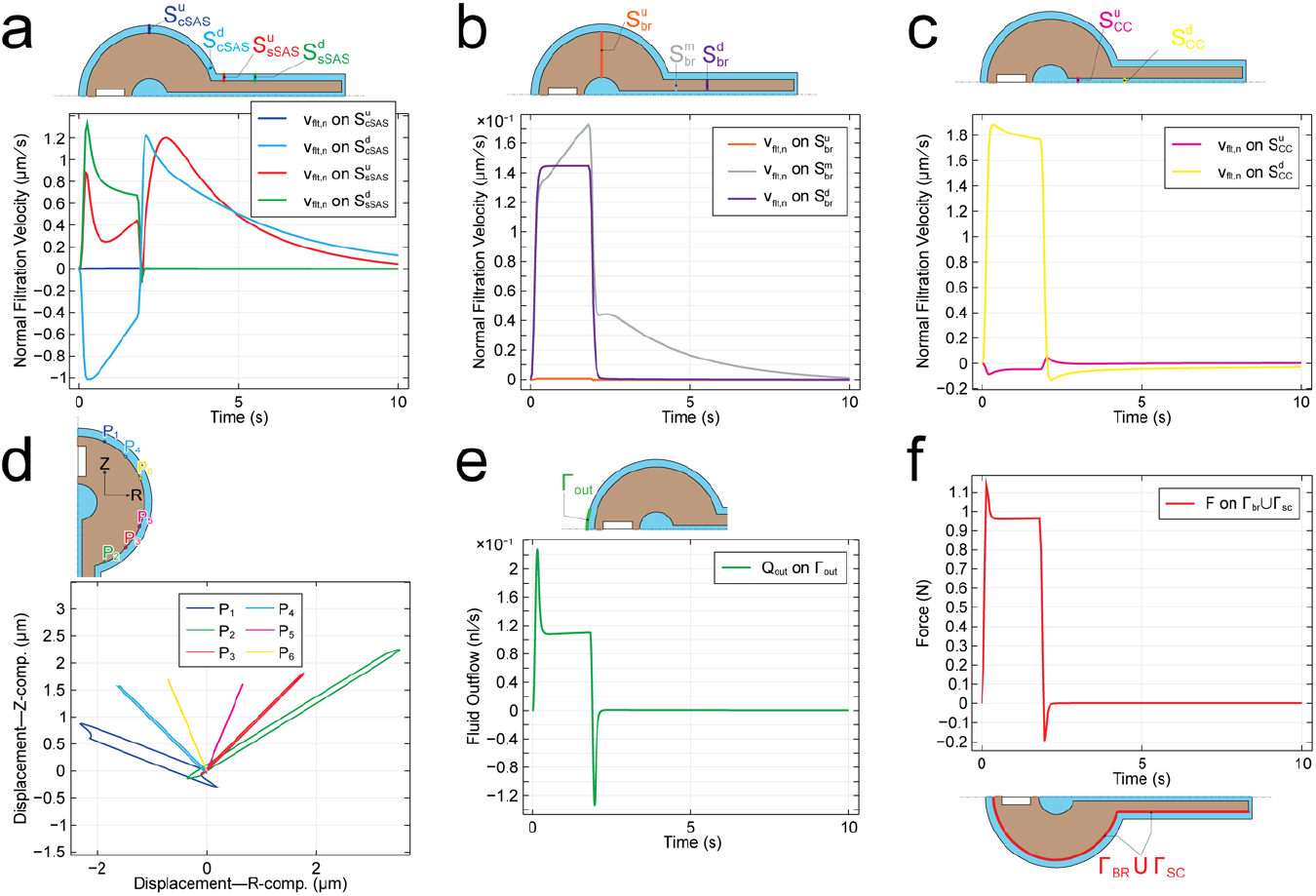
Computational results from a finite element simulation of the flow induced by a squeeze of intensity *p*_0_ = 20mmHg applied over the SZ. The duration of the squeeze pulse is 2s. The duration of the simulation is 10s. The simulation is based on Equations (1)—(9). The boundary conditions are described in the Supplementary Material. The parameters used in the simulation are found in Supplementary Table 1. **Note: the resistance scaling factors adopted here are** α_cs_ = 10^6^ **and** α_out_ = 6 × 10^8^. **a**. Average of normal filtration velocity (in μm/s) over each of the cranial and spinal SAS sections (shown in the inset) over time. The plot displays 4 lines, with the blue one appearing as horizontal line near zero –due to the different orders of magnitude of the filtration velocity across the different SAS sections. The unit normal vector to the sections points in the rostral direction. **b**. Average of normal filtration velocity (in μm/s) over each of the brain and spinal cord sections (shown in the inset) over time. The plot displays 3 lines, with the orange one appearing as horizontal line near zero –due to the different orders of magnitude of the filtration velocity across Ω_BR_. The unit normal vector to the sections points in the rostral direction. **c**. Average of normal filtration velocity (in μm/s) over each of the central canal sections (shown in the inset) over time. The unit normal vector to the sections points in the rostral direction. **d**. Trajectories of points P1–P6 (shown in the inset) on the surface of the brain: traces of the points indicated in the inset over the time interval 0 < *t* < 10 s. **e**. Volumetric fluid outflow Q_out_ (in n**F**/s) through the outlet boundary Γ_out_ over time. Q_out_ > 0: fluid flow out of Ω_SAS_. Q_out_ is computed as the integral of the normal component of filtration velocity over the surface indicated. **f**. Average force *F* (in N) exerted by CSF over time onto brain and spinal cord during the squeeze. *F*(*t*) is computed as the integral average of (*m ·* T*m*) over the surface Γ_bζ_ ∪ Γ_sc_, where T is the total Cauchy stress acting on the mixture in the SAS and *m* is the outward unit normal to the surface indicated.

**Supplementary Figure 12.**
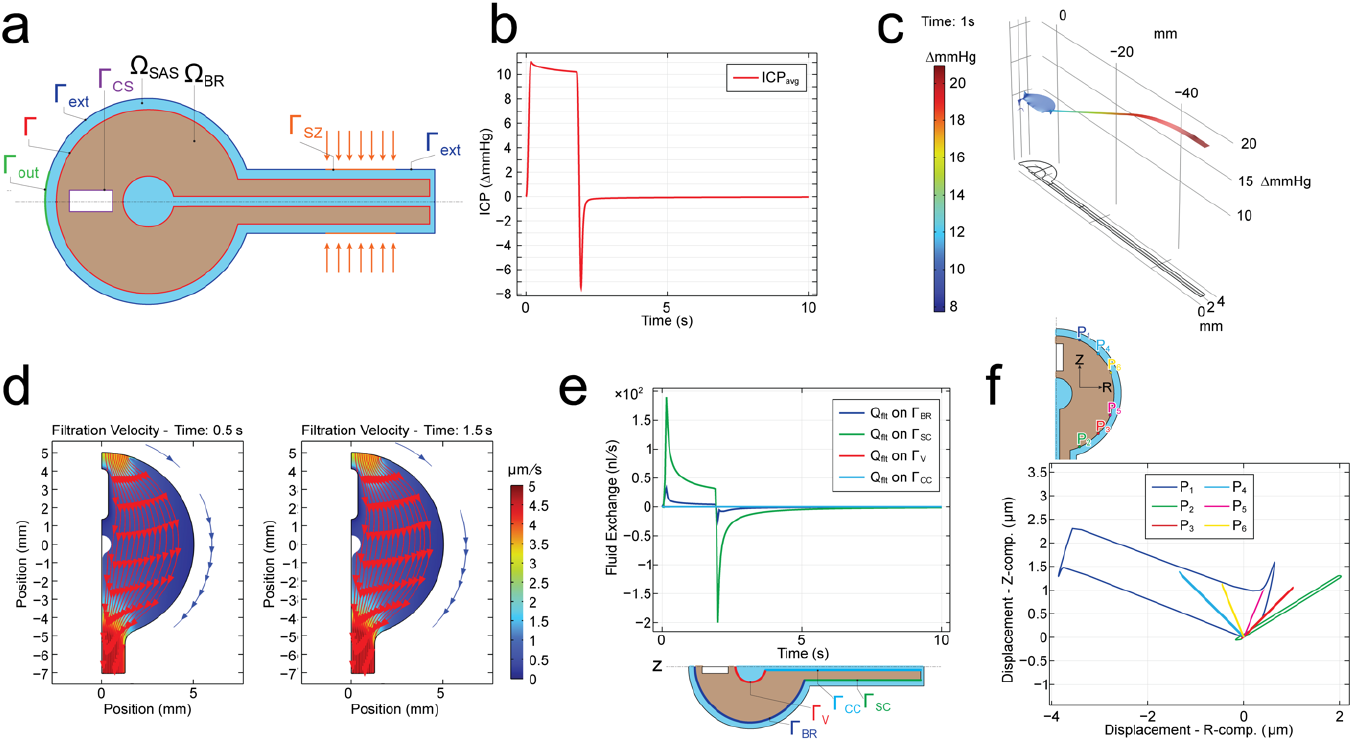
Computational results from a finite element simulation of the flow induced by a squeeze of intensity *p*_0_ = 20mmHg applied over the SZ. The duration of the squeeze pulse is 2s. The duration of the simulation is 10s. The simulation is based on Equations (1)—(9). The boundary conditions are described in the Supplementary Material. The parameters used in the simulation are found in Supplementary Table 1. **Note: the resistance scaling factors adopted here are** α_cs_ = 10^10^ **and** α_out_ = 6 × 10^4^. **a**. Initial geometry (not to scale) detailing model domains and boundaries. Ω_BR_: brain and spinal cord domain (pale pink); Ω_SAS_: CSF-filled domain (cyan); Γ: Ω_BR_ − Ω_SAS_ interface (red); Γ_ext_: external boundary of meningeal layer (blue); Γ_sZ_: squeeze zone (orange); Γ_out_: outlet boundary representing the cribriform plate CSF outflow pathway (green); Γ_cs_: central sinus boundary (purple). **b**. Average of pore pressure (in mmHg) over Ω_BR_ excluding the spinal cord over time. **c**. Spatial distribution of pore pressure (in mmHg) over Ω_BR_ ∪ Ω_SAS_ at *t* = 1 s during the squeeze pulse. **d**. Streamlines of filtration velocity ***ν***_flt_ (i.e., curves tangent to filtration velocity field; red arrows) within Ω_BR_ excluding the spinal cord, at *t* = 0.5 s (left) and *t* = 1.5 s (right) during the squeeze pulse, overlaying the color plot of the filtration velocity magnitude (in μm/s), computed as 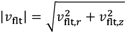. Because the SAS is extremely thin, it is not meaningful to show a full plot of the streamlines in the SAS. This said, the blue line with arrows placed on the right side of each streamline plot is meant to indicate the direction of flow in the SAS at the corresponding time. **e**. Volumetric fluid exchange rate Q_flt_ (in n**F**/s) over time across: the brain shell surface Γ_bζ_ (blue), spinal cord surface Γ_sc_ (green), ventricle surface Γ_***x***_ (red), and central canal surface Γ_cc_ (light blue). Q_flt_ > 0: fluid flow from Ω_BR_ into Ω_SAS_. Q_flt_ is computed as the integral of the normal component of filtration velocity over the surface indicated. The plot displays 4 lines, two that are easily seen (blue and green lines), and two that overlap and appear as horizontal lines near zero (red and light blue lines). This is due to the different orders of magnitude of Q_flt_ across the different portions of Γ. **f**. Trajectories of points P1–P6 (shown in the inset) on the surface of the brain: traces of the points indicated in the inset over the time interval 0 < *t* < 10 s.

**Supplementary Figure 13.**
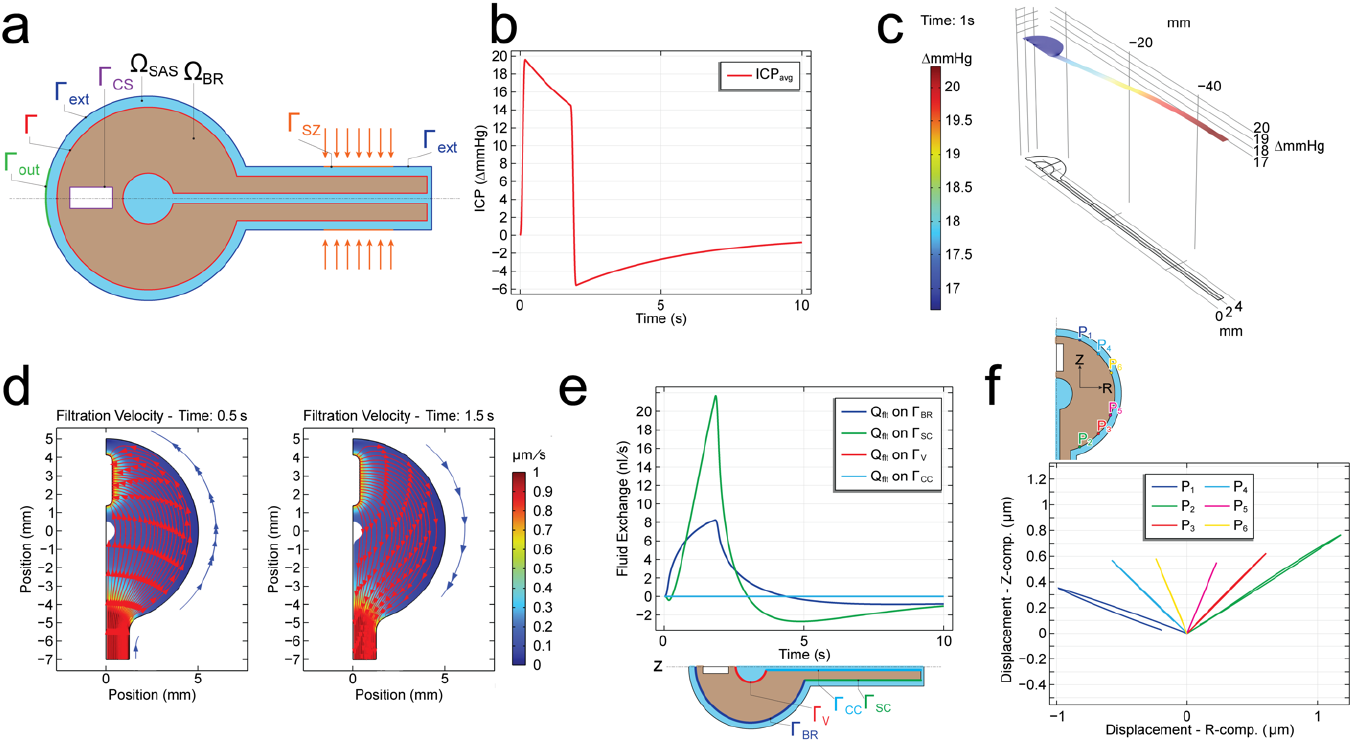
Computational results from a finite element simulation of the flow induced by a squeeze of intensity *p*_0_ = 20mmHg applied over the SZ. The duration of the squeeze pulse is 2s. The duration of the simulation is 10s. The simulation is based on Equations (1)—(9). The boundary conditions are described in the Supplementary Material. The parameters used in the simulation are found in Supplementary Table 1. **Note: the resistance scaling factors adopted here are** α_cs_ = 10^8^ **and** α_out_ = 6 × 10^8^. **a**. Initial geometry (not to scale) detailing model domains and boundaries. Ω_BR_: brain and spinal cord domain (pale pink); Ω_SAS_: CSF-filled domain (cyan); Γ: Ω_BR_ − Ω_SAS_ interface (red); Γ_ext_: external boundary of meningeal layer (blue); Γ_sZ_: squeeze zone (orange); Γ_out_: outlet boundary representing the cribriform plate CSF outflow pathway (green); Γ_cs_: central sinus boundary (purple). **b**. Average of pore pressure (in mmHg) over Ω_BR_ excluding the spinal cord over time. **c**. Spatial distribution of pore pressure (in mmHg) over Ω_BR_ ∪ Ω_SAS_ at *t* = 1 s during the squeeze pulse. **d**. Streamlines of filtration velocity ***ν***_flt_ (i.e., curves tangent to filtration velocity field; red arrows) within Ω_BR_ excluding the spinal cord, at *t* = 0.5 s (left) and *t* = 1.5 s (right) during the squeeze pulse, overlaying the color plot of the filtration velocity magnitude (in μm/s), computed as 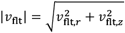. Because the SAS is extremely thin, it is not meaningful to show a full plot of the streamlines in the SAS. This said, the blue line with arrows placed on the right side of each streamline plot is meant to indicate the direction of flow in the SAS at the corresponding time. **e**. Volumetric fluid exchange rate Q_flt_ (in n**F**/s) over time across: the brain shell surface Γ_bζ_ (blue), spinal cord surface Γ_sc_ (green), ventricle surface Γ_***x***_ (red), and central canal surface Γ_cc_ (light blue). Q_flt_ > 0: fluid flow from Ω_BR_ into Ω_SAS_. Q_flt_ is computed as the integral of the normal component of filtration velocity over the surface indicated. The plot displays 4 lines, two that are easily seen (blue and green lines), and two that overlap and appear as horizontal lines near zero (red and light blue lines). This is due to the different orders of magnitude of Q_flt_ across the different portions of Γ. **f**. Trajectories of points P1–P6 (shown in the inset) on the surface of the brain: traces of the points indicated in the inset over the time interval 0 < *t* < 10 s.

**Supplementary Figure 14.**
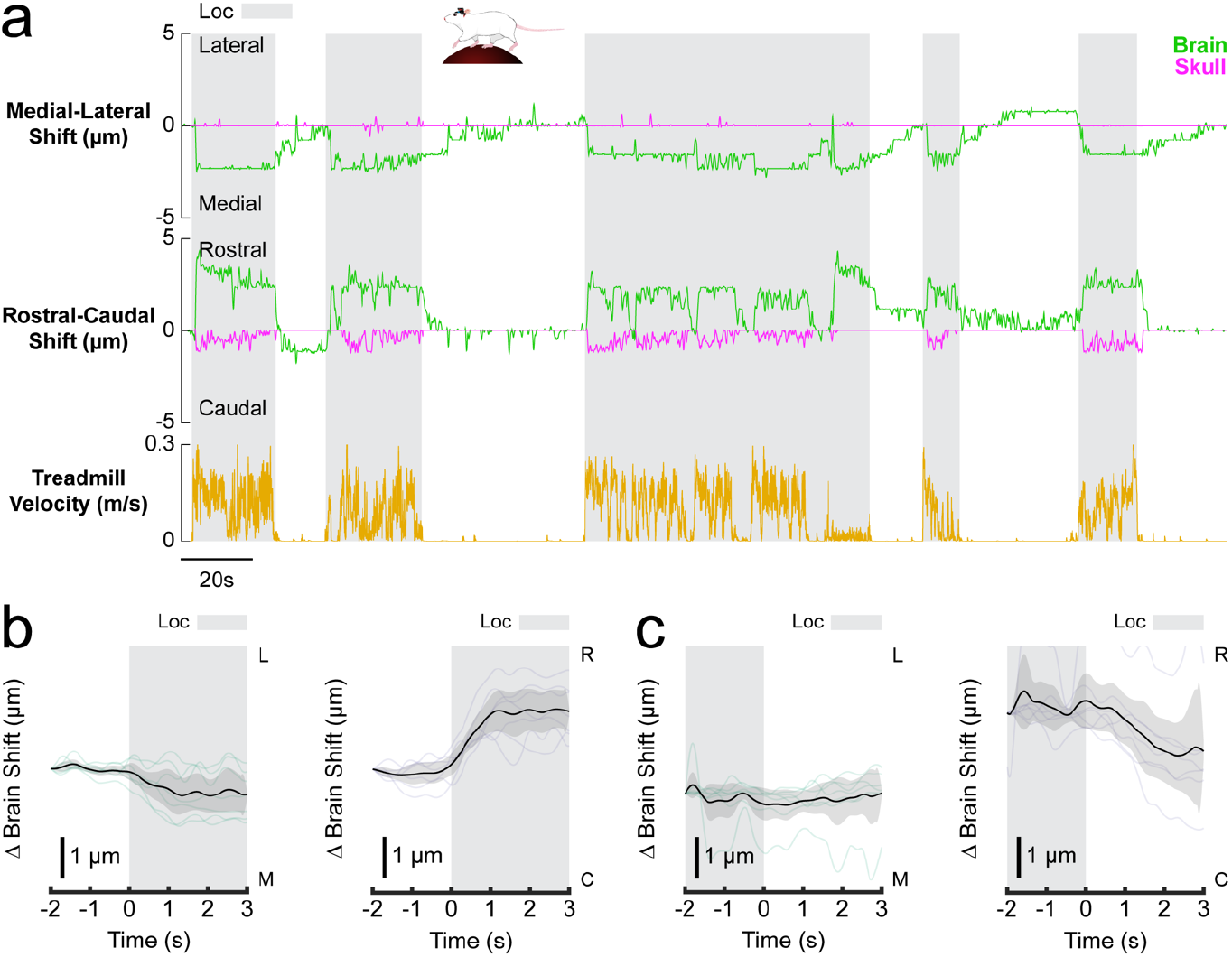
Motion of the olfactory bulb was rostral and medial. **a**. A single trial from an olfactory bulb showing brain (green) and skull (magenta) motion as well as locomotion (black). Like the cortex, locomotion events resulted in rostral motion of the olfactory bulb. However, the olfactory bulbs exhibited medial displacement instead of lateral. **b**. Locomotion-triggered average olfactory bulb motion for each trial with the average of these plotted in black with the 90 percent confidence interval. **c**. Locomotion cessation-triggered average olfactory bulb motion for each trial with the average of these plotted in black with the 90 percent confidence interval.

**Supplementary Figure 15.**
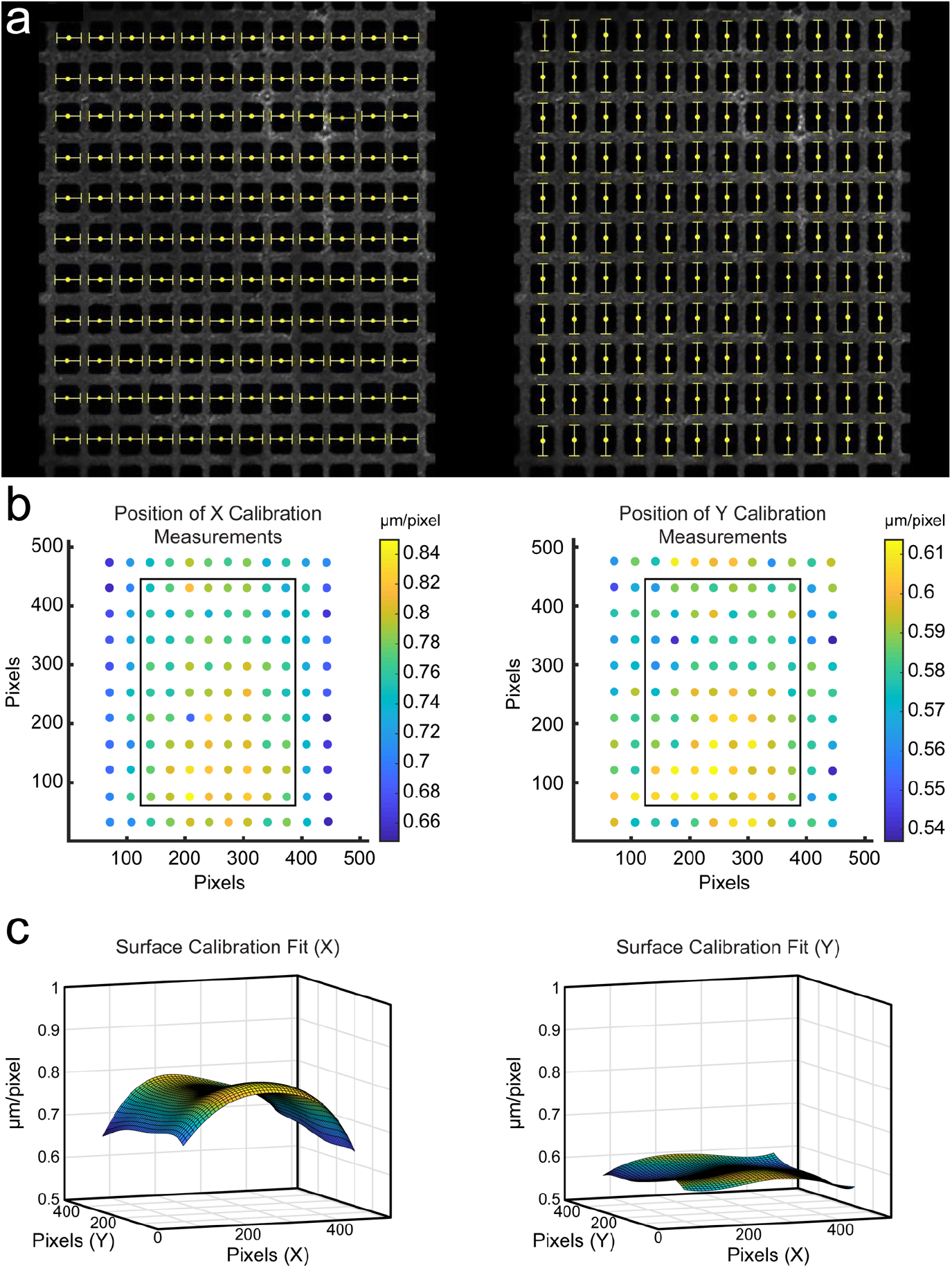
Imaging calibration. **a**. Image of the copper mesh used for calibration. Each 19µm x 19µm hole had its height and width measured in pixels using a full width at half maximum algorithm (yellow bars). **b**. Pixels per micron in the X (left) and Y (right) dimensions. Box shows imaged area. **c**. Fitted surfaces for X (left) and Y (right).

**Supplementary Figure 16.**
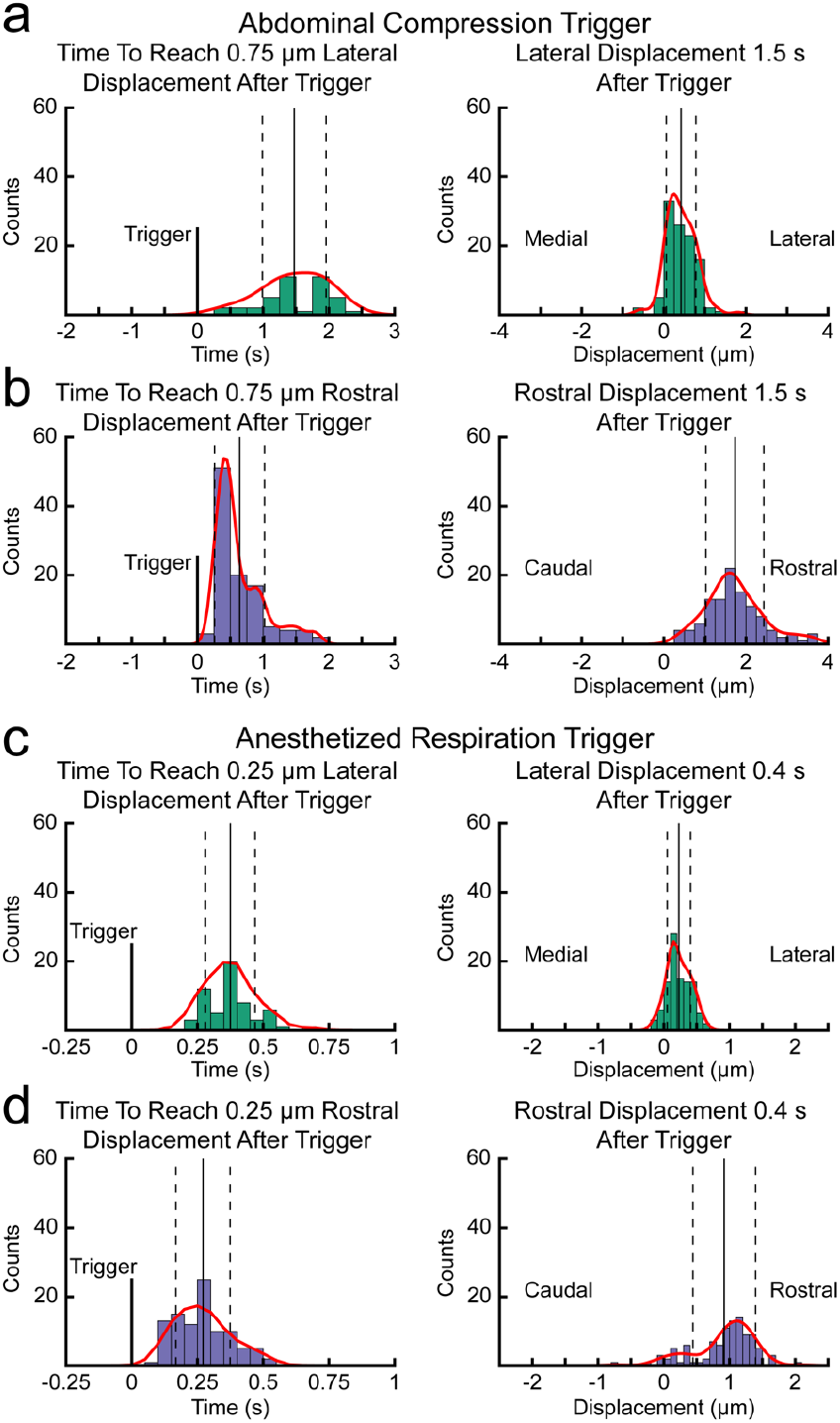
Brain displacement speed during anesthesia and externally applied abdominal pressure. **a**. Histogram of the amount of time it takes the brain to displace laterally 0.75 µm following the onset of an abdominal compression with the probability density function, mean and standard deviation (left). Histogram of lateral displacement of the brain 1.5 seconds following the onset of an abdominal compression with the probability density function, mean and standard deviation (right). **b**. Histogram of the amount of time it takes the brain to rostrally displace 0.75 µm following the onset of an abdominal compression with the probability density function, mean and standard deviation (left). Histogram of rostral displacement of the brain 1.5 seconds following the onset of an abdominal compression with the probability density function, mean and standard deviation (right). **c**. Histogram of the amount of time it takes the brain to displace laterally 0.75 µm following the onset of an anesthetized respiration event with the probability density function, mean and standard deviation (left). Histogram of the lateral displacement of the brain 1.5 seconds following the onset of an anesthetized respiration event with the probability density function, mean and standard deviation (right). **d**. Histogram of the amount of time it takes the brain to rostrally displace 0.75µm following the onset of an anesthetized respiration event with the probability density function, mean and standard deviation (left). Histogram of the rostral displacement of the brain 1.5 seconds following the onset of an anesthetized respiration event with the probability density function, mean and standard deviation (right).

**Supplementary Figure 17.**
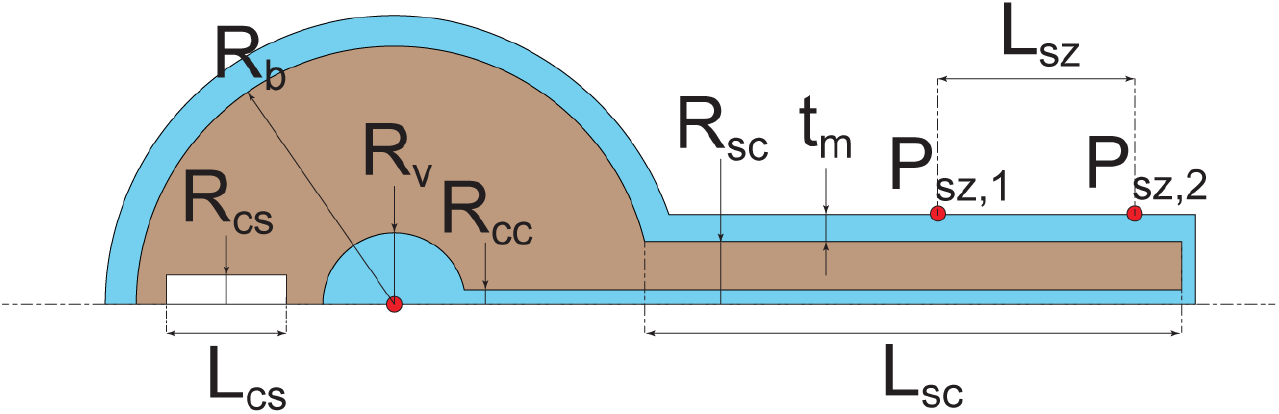
Initial geometry (not to scale) of an axially symmetric poroelastic domain (pale pink) representing brain and spinal cord immersed in CSF-filled poroelastic compartment (cyan), including the cranial and spinal SAS, central canal and a spherical ventricle representing the ventricular system. All geometric parameters defining the model, together with the chosen values are summarized in Table 1. In the geometry implemented in COMSOL Multiphysics for all finite element simulations, corners are rounded using a “fillet” tool.

**Movie 1. Brain motion is rigid**. A single-plane video of a mouse brain through a cranial window was tracked in eight locations. The high degree of similarity between the calculated movement at each location demonstrates the robustness of the template-matching tracking program, the accuracy of the two-dimensional distortion calibration, and a rigid shift of the parenchyma within the imaging plane.

**Movie 2. Relationship of brain motion to abdominal muscle EMG activity, locomotion, and respiration**. The brain moves rostro-laterally in response to abdominal muscle contractions prior to and during locomotion events. Respiration does not drive brain motion during the resting phase in the awake state. Two and three-dimensional figures are included to demonstrate the cranial window environment used to capture the brain and skull motion *in vivo*.

**Movie 3. Brain motion without locomotion (hunching)**. Prior to locomotion, the mouse exhibits a hunching behavior that changes its posture and invokes abdominal muscle contraction. This results in rostro-lateral motion of the brain without the presence of locomotion activity. Shortly after, the mouse begins a locomotion event that shows a higher degree of abdominal muscle contraction and increased rostro-lateral brain motion. These data show that while locomotion events can predict motion of the brain within the skull, it is not required for brain motion.

**Movie 4. Simultaneous skull, dura, and brain tracking**. The electrically-tunable lens was programmed for simultaneous capture of three layers to track skull, dura, and brain motion. As seen in the three-dimensional reconstruction of the cranial window, the dural vessel (white) is much closer to the parenchyma surface (green) than the fluorescent microspheres on the window (magenta) as it resides on the internal surface of the skull. The data demonstrate that when the brain moves, the dura remains stationary with the skull despite the small size of the subarachnoid space.

**Movie 5. Respiration-linked brain motion during anesthesia**. Respiration-driven brain motion was not observed during the awake and behaving state in mice. However, brain motion was occasionally detected during periods of deep anesthesia. In this example, the mouse exhibits very little brain motion when anesthetized with 1 percent isoflurane in oxygen in the first 20 seconds of data collection. The isoflurane was then increased to 5 percent in oxygen to generate deeper and slower respiration. In this state, the abdominal muscles are more strongly recruited in each breath and the brain exhibits a rostro-lateral shift within the skull. Reducing the isoflurane to 1 percent in oxygen at the end of the data set resulted in reduced abdominal muscle contraction force and less brain motion. These results suggest that brain motion can only be driven by respiration in mice when the abdominal muscle contractions associated with each breath generate sufficient pressure changes within the abdomen.

**Movie 6. MicroCT of vertebrae and vertebral venous plexus**. This three-dimensional segmentation of a mouse microCT shows how vasculature inside and outside of the vertebral bones are oriented. Furthermore, it demonstrates how the vessels connect through the ventral surface of the individual vertebrae. The bone transitions between opaque and transparent to display the entirety of the vertebral venous plexus.

**Movie 7. Brain motion induced by abdominal compression**. A pressure cuff wrapped around the abdomen of a lightly anesthetized mouse was used to induce an increase in intra-abdominal pressure for two seconds. These pressure increases resulted in rostro-lateral motion of the brain in the skull for the duration of the compression and a return to baseline position following pressure release. These results suggest that externally applied intra-abdominal pressure changes can drive brain motion when controlling for behavior in a mouse model.

**Movie 8. Olfactory bulb motion**. The olfactory bulb moves rostrally within its skull compartment during locomotion, similar to the cortex. However, the olfactory bulbs shift medially as well, in contrast to the lateral motion seen in the cortex. Following locomotion, the olfactory bulb begins to move laterally to return to its baseline position but also overshoots its resting position caudally before slowly returning rostrally. This movement behavior is unique to the olfactory bulbs and suggests a difference in brain motion mechanics between the olfactory bulbs and the cortical hemispheres.

**Animation 1. Gut-brain hydraulic axis**. Reducing the volume of the abdominal cavity increases intra-abdominal pressure, forcing blood into the vertebral venous plexus. This narrows the dural sac and forces cranial cerebrospinal fluid flow, resulting in increased intracranial pressure and brain motion within the skull.

## Acknowledgements

We thank Micheal Tribone for mouse illustrations, and Sutter Instruments for assistance in design and machining the ETL mount.

